# Auxin-induced nanoclustering of membrane signaling complexes underlies cell polarity establishment in Arabidopsis

**DOI:** 10.1101/734665

**Authors:** Xue Pan, Linjing Fang, Jianfeng Liu, Betul Senay-Aras, Wenwei Lin, Shuan Zheng, Tong Zhang, Uri Manor, Weitao Chen, Zhenbiao Yang

## Abstract

Cell polarity is fundamental to the development of both eukaryotic and prokaryotic organisms, yet the mechanism of its establishment remains poorly understood. Here we show that signal-activated nanoclustering of membrane proteins and a cytoskeleton-based feedback loop provide an important mechanism for the establishment of cell polarity. The phytohormone auxin promoted sterol-dependent nanoclustering of cell surface transmembrane receptor-like kinase 1 (TMK1) to initiate cell polarity during the morphogenesis of Arabidopsis puzzle piece-shaped leaf pavement cells (PC). Auxin-triggered nanoclustering of TMK1 stabilized flotillin-associated ordered nanodomains, which were essential for auxin-mediated formation of ROP6 GTPase nanoclusters that act downstream TMK1 to promote cortical microtubule ordering. Mathematical modeling further demonstrated the essential role of this auxin-mediated stabilization of TMK1 and ROP6 nanoclusters, and predicted the additional requirement of ROP6-dependent cortical microtubules for further stabilization of TMK1-sterol nanodomains and the polarization of PC. This prediction was experimentally validated by genetic and biochemical data. Our studies reveal a new paradigm for polarity establishment: A diffusive signal triggers cell polarization by activating cell surface receptor-mediated lateral segregation of signaling components and a cytoskeleton-mediated positive feedback loop of nanodomain stabilization.

**Highlights:**

- Sterols are required for cell polarity in Arabidopsis leaf epidermal cells
- Auxin promotes lipid ordering and polar distribution of ordered lipid nanodomains at the plasma membrane (PM)
- Auxin stabilizes sterol-dependent nanoclustering of transmembrane kinase (TMK1), a PM auxin signal transducer
- Auxin-induced TMK1 nanoclustering is required but insufficient for cell polarization
- Microtubule-based feedback stabilization of the auxin-induced TMK1 nanodomains can generate cell polarity

## Introduction

Cell polarity is a fundamental cellular property required for the specialization and function of essentially all cells. The initiation of cell polarity requires spatially controlled cell signaling and the dynamic spatiotemporal organization of signaling molecules at the cell surface (Etienne-Manneville and Hall, 2002; Johnson, 1999; Yang and Lavagi, 2012). How this spatiotemporal organization is initiated, especially by uniform signals, remains unclear despite decades of studies into the mechanisms behind the establishment and maintenance of cell polarity. To date two prevalent models have been proposed to explain self-organizing cellular symmetry breaking or that induced by spatial cues: asymmetric recruitment of polarity signaling proteins or endomembrane-linked trafficking of these proteins (Butler and Wallingford, 2017; Nelson, 2003; Strutt and Strutt, 2009; Vladar et al., 2009). However, these models have difficulty explaining how a uniform field of highly diffusible small signaling molecules can lead to symmetry breaking. Furthermore, neither of these models consider the heterogeneity of the plasma membrane (PM) lipid environment. Increasing evidence shows that many cell surface-signaling proteins are not uniformly mixed with membrane lipids via free-diffusion based equilibrium, but form nanoclusters with specialized lipids (Garcia-Parajo et al., 2014; Harding and Hancock, 2008; Platre et al., 2019; Sezgin et al., 2015; Sezgin et al., 2017; Zhou and Hancock, 2015). Membrane domains enriched in sterols or saturated lipids, often referred to as lipid rafts, have long been suggested as a crucial platform for receptor-mediated signal transduction at the cell surface, particularly in the spatial regulation of key cellular processes, such as cell polarization (Gomez-Mouton et al., 2001; Makushok et al., 2016; Schuck and Simons, 2004; Simons, 2018; Simons and Sampaio, 2011) and immune response (Stone et al., 2017; Varshney et al., 2016; Vieira et al., 2010). In addition to sterol-dependent nanodomains, anionic lipids have been implicated in the regulation of nanoclusters of small GTPases, such as Ras, Cdc42 and ROP6 (a Rho-like GTPase from plants), on the PM (Platre et al., 2019; Sartorel et al., 2018; Zhou et al., 2017; Zhou et al., 2015). Spatial segregation of signaling molecules into nanoclusters at the PM is a generic phenomenon during signal transduction (Ariotti et al., 2010; Garcia-Parajo et al., 2014; Harding and Hancock, 2008). However, it remains unclear how distinct nanocluster-based signaling platforms are established and maintained by specific signals and how they may contribute to the spatiotemporal regulation of cellular processes. Here we show that in Arabidopsis the highly diffusible small molecule auxin regulates the formation of distinct PM nanoclusters via its PM signaling pathway, leading to cell polarization.

Auxin, a molecule tightly linked with polarity (Pan et al., 2015), impacts virtually every aspect of plant growth and development via nuclear gene transcription and rapid non-transcriptional responses controlled by the nuclear/cytoplasmic TIR1/AFB auxin receptors (Chapman and Estelle, 2009; Dindas et al., 2018; Fendrych et al., 2018; Leyser, 2018; Monshausen et al., 2011; Salehin et al., 2015; Scheitz et al., 2013), as well as those mediated by the PM-localized transmembrane kinases (TMKs) (Cao et al., 2019; Xu et al., 2010). Auxin-induced activation of ROPs is involved in cellular symmetry breaking such as polarization of PIN proteins, which export auxin (Nagawa et al., 2012; Xu et al., 2010), and the morphogenesis of *Arabidopsis thaliana* leaf and cotyledon pavement cells (PCs) (Chen et al., 2015; Fu et al., 2005; Xu et al., 2014; Xu et al., 2010). These cells form the puzzle-piece shape with interlocking lobes and indentations, which require the establishment of multiple alternating polar sites for the formation of lobes and indentations, respectively, herein we term multi-polarity (Fig. 1A). In PCs, the multi-polarity formation requires the TMK-dependent activation of ROPs by auxin (Xu et al., 2014). In particular, ROP6 is polarly localized to and defines the indentation regions where it promotes the ordering of cortical microtubules (CMT) (Fu et al., 2009; Lin et al., 2013). In this study, we found that auxin promotes the formation of inter-dependent cell surface TMK1 nanoclusters and ordered lipid nanodomains as well as microtubule-mediated diffusion restriction of these TMK1/ordered lipid nanodomains to activate symmetry breaking for PC multi-polarity formation.

**Fig 1.**
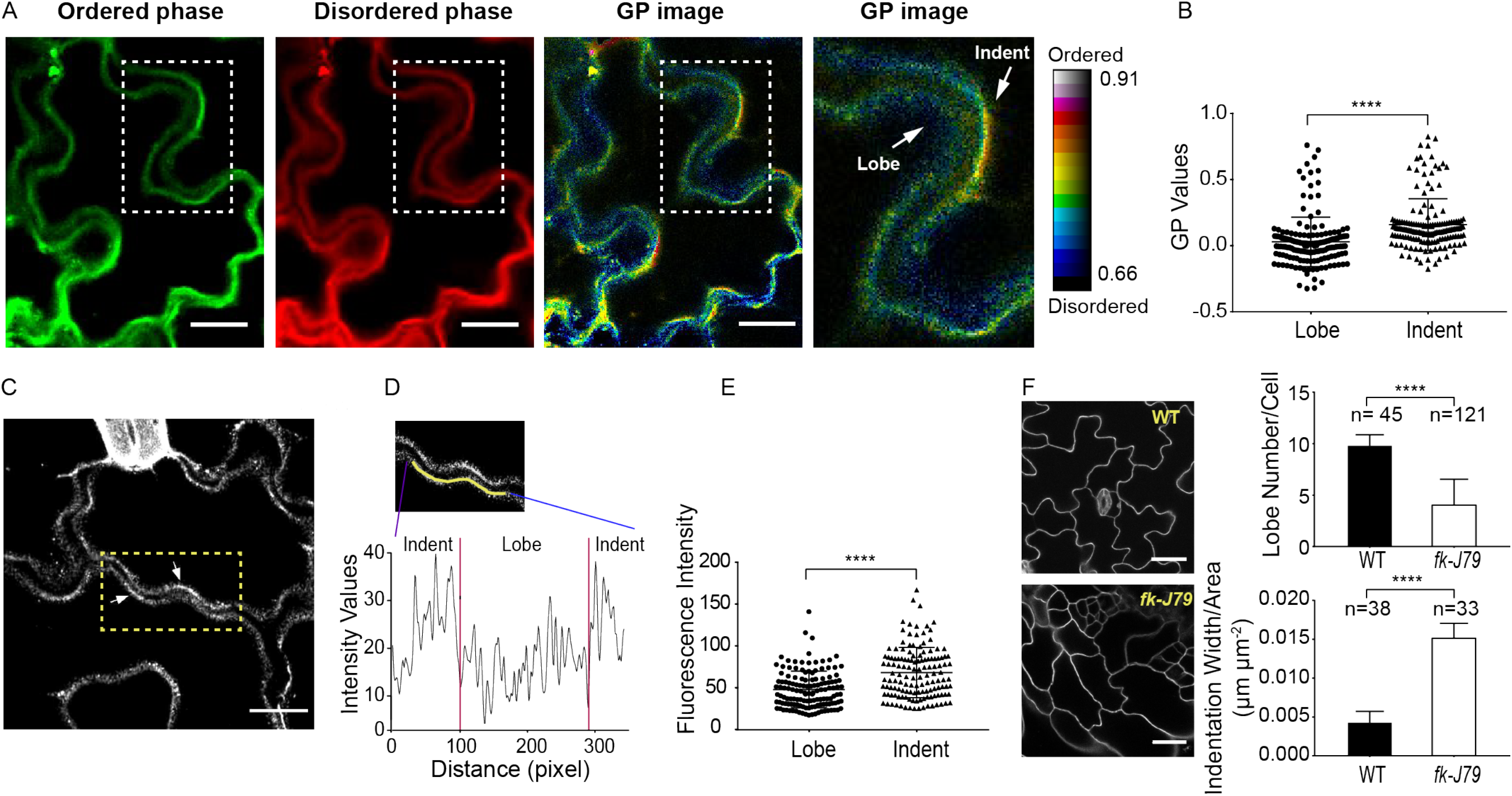
High lipid-order membrane domains have an indenting region-preferred localization. **(A,B)** Plasma membrane order visualization using di-4-ANEPPDHQ staining in pavement cells of 2-3 days-old cotyledons. **(A)** Representative images obtained after di-4-ANEPPDHQ staining. **1^st^ panel:** di-4-ANEPPDHQ fluorescence recorded between 500-580 nm, representing high lipid order. **2^nd^ panel:** di-4-ANEPPDHQ fluorescence recorded between 620-750nm, representing low lipid order. **3^rd^ panel:** radiometric color-coded GP image generated from 1^st^ and 2^nd^ panels. **4^th^ panel:** the enlarged radiometric color-coded GP images generated from the boxed areas shown in the left three panels. GP image is a false-color image, which runs over the range indicated by the color bar. The colorbar values representing the GP values ascend from bottom to top, with red colors indicating high membrane order, whereas blue colors indicating low membrane order. Scale bars = 15 μm. **(B)** Quantitative analysis of mean GP values obtained from the complementary lobing and indenting regions of 161 sites of 56 cells. GP values at indenting regions are higher than that at lobing regions, indicating higher membrane lipid order in indenting regions. **(C,D,E)** Flot1∷Flot1-mVenus shows a polar distribution towards the indenting regions. **(D)** Representative image showing the distribution of Flot-mVenus in PCs of 2-3 days-old cotyledons. Scale bars = 15 μm. **(D)** Fluorescent intensity values scanned along the line indicated in the top panel. **(E)** Quantitative analysis of fluorescence intensity at the complementary lobing and indenting regions of 138 sites of 45 cells. **(F)** The sterol biosynthesis mutant fackel-J79 (*fk-J79)* shows altered pavement cell shape. The *fk-J79* mutant is defective in gene encoding a sterol C14 reductase. Homozygous *fk-J79* seedlings have an altered sterol composition, with reduced levels of campesterol and sitosterol but increased stigmasterol compared with wild type. **Left panels**: Representative images of pavement cells in cotyledons of *fk-J79* mutant (bottom panel) and its corresponding wild-type (top panel). Scale bars = 30 μm. **Right panels:** Quantitative analysis of the number of lobes (top panel) and indentation widths (bottom panel) for *fk-J79* mutant and its wild-type. The *fk-J79* mutant has decreased lobe number per cell and increased indentation width compared to its wild type. *n* represents the number of cells. Similar phenotypic changes were observed for other sterol biosynthesis mutants (See supplementary fig. S2). Data were presented as mean ± SD. See methods for detailed statistical analyses.

## Results

### Ordered lipid nanodomains are polarized and critical for multi-polarity formation in leaf pavement cells

To assess the role of lipid ordering in the morphogenesis of the puzzle-piece shaped PCs, we analyzed the distribution of ordered lipid domains in cotyledon PCs using di-4-ANEPPDHQ, a lipid order-sensitive probe (Gerbeau-Pissot et al., 2016; Jin et al., 2005; Owen et al., 2012; Zhao et al., 2015). Based on the fluorescent intensity of the probe in two different spectral channels, green (ordered) and red (disordered), a ratiometric general polarized image (GP image) was generated. High GP value (increased green fluorescence compared with red) represents the highly ordered membrane lipids. Plasmolysis with 0.8 M mannitol revealed that the peripheral staining is associated with the plasma membrane, but not with the cell wall (Fig. S1). The GP images showed that the localization of ordered lipid domains is biased towards the indentation region (Fig. 1A). Further quantitative analysis revealed that GP values extracted from indentation regions are much higher than those extracted from lobing regions (Fig. 1B). To complement this staining method, the localization of flotillin-1, a well-established marker for ordered sterol-rich nanodomains in both plant and mammalian systems (Glebov et al., 2006; Li et al., 2012), was visualized. Consistent with previous studies, flotillin proteins form dynamic nanoclusters in the PM (Supplementary Video 1 and Video 2). Our analyses further demonstrated that in the majority of cases, flotillin proteins were more abundant in indentations than in lobes (Fig. 1C-E, Supplemental Video 3). Collectively, these results show that ordered lipid domains are preferentially distributed to the indentation region of the PC plasma membrane, as is the active ROP6 GTPase (Fu et al., 2009).

To evaluate the roles of sterols in the formation of the polarity in the in PCs, we examined PC phenotypes of sterol biosynthesis mutants, which were previously reported to have altered sterol composition (Jang et al., 2000; Men et al., 2008; Schrick et al., 2000; Souter et al., 2002) and auxin signaling (Men et al., 2008; Souter et al., 2002). All these mutants display significantly altered PC shape with reduced lobe number and increased indentation width compared to their wild-type control (Fig. 1F and Fig. S2). The depletion of sterol from the PM by methyl-β-cyclodextrin (mβCD) (Kierszniowska et al., 2009; Mahammad and Parmryd, 2015), a sterol-depleting agent, also caused a similar defect in PC morphogenesis (see below). These defects resemble those induced by the disruption of the ROP6 pathway that promotes the organization of cortical microtubules and the formation of PC indentation (Fu et al., 2005; Fu et al., 2009). The PC phenotypes induced by sterol-deficiency and the distribution of ordered lipid domains to the indentation regions suggest sterol-rich ordered lipid domains, together with ROP6, polarly define the indentation regions, herein termed the indentation polarity, and are critical for the formation of the indentation polarity.

### Auxin promotes lipid ordering required for the indentation polarity

Our previous studies show that auxin promotes the formation of the puzzle-piece PC shape (Xu et al., 2014; Xu et al., 2010), suggesting that auxin is a signal that initiates the establishment of the multi-polarity. Thus, we asked whether auxin regulates the generation of ordered lipid domains in PCs. The auxin biosynthesis mutant *wei8-1tar2-1* with reduced auxin levels in cotyledons (Stepanova et al., 2008) exhibited greatly reduced lipid order in PCs, as indicated by di-4-ANEPPDHQ staining (Figs. 2A and 2B). This defect in lipid ordering was rescued by exogenous auxin (Figs. 2A and 2B). Similar to the *wei8-1tar2-1* mutant, the sterol biosynthesis mutant *fk-J79* also exhibited reduced lipid order (Figs. 2C and 2D). However, unlike the *wei8-1tar2-1* mutant, the *fk-J79* mutant was completely insensitive to exogenous auxin in the promotion of lipid ordering (Figs. 2C-2D). Furthermore, auxin promotion of PC multi-polarity was completely abolished in the *fk-J79* mutant (Fig. 2E).

**Fig 2.**
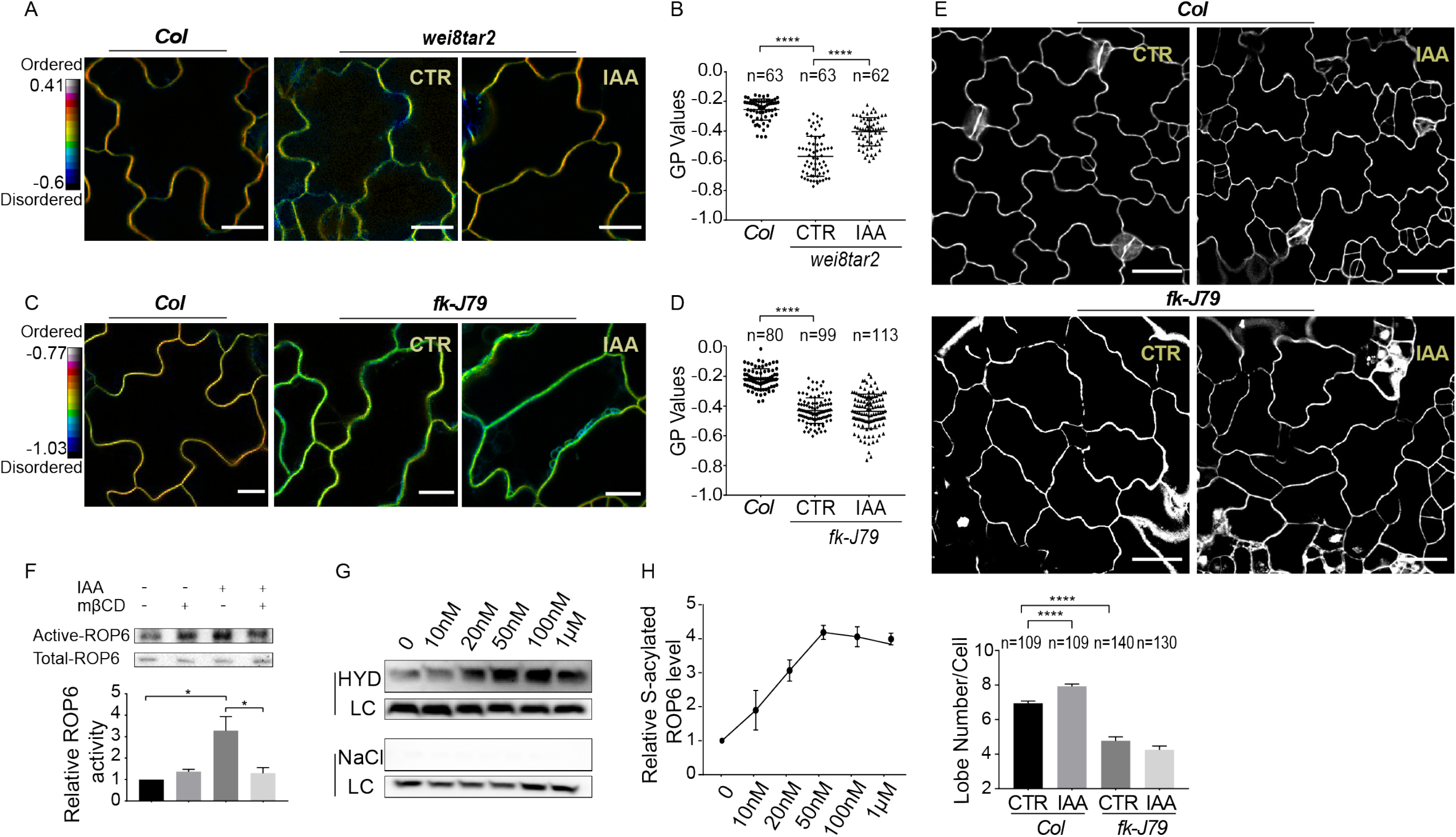
Auxin promotes lipid ordering required for ROP6 activation. **(A,B)** The auxin biosynthesis mutant *wei8-1tar2-1* has reduced plasma membrane order (green to yellow) compared to the wild-type (red to orange). Adding 50nM of IAA for 3hrs rescued the reduced membrane order phenotype of *wei8-1tar2-1* mutant. **(A)** Representative GP images of pavement cells in wild-type (**left panel**) and *wei8-1tar2-1* mutant without (**middle panel**) and with IAA treatment (**right panel**) obtained after di-4-ANEPPDHQ staining. Scale bars = 15 μm. **(B)** Quantitative analysis of mean GP value extracted from the plasma membrane of multiple pavement cells. (**C,D**) Plasma membrane order is dramatically reduced in *fk-J*79 mutant (from green to yellow) compared to wild-type (from red to orange). The reduced lipid order in *fk-J79* mutant is not rescued by the exogenously supplied IAA. **(C)** Representative GP images of pavement cells in wild-type (**left panel**) and *fk-J*79 mutant without (**middle panel**) and with IAA treatment (**right panel**) obtained after di-4-ANEPPDHQ staining. Scale bars = 15 μm. **(D)** Quantitative analysis of mean GP value extracted from the plasma membrane of multiple pavement cells. **(E)** Auxin-promoted pavement cell interdigitation is abolished in the *fk-J*79 mutant. **(E, top panels)** Representative confocal images of pavement cells in wild-type (Col-0) and *fk-J79* with and without the IAA treatment. Scale bars = 30 μm. **(E, lower panel)** The average lobe number per cell increased in wild-type after IAA treatment, but did not change in *fk-J*79 mutant. Data were presented as mean ± SEM. See methods for detailed statistical analyses. **(F) Top panel:** Representative Western blot image showing that deprivation of membrane sterols by methyl-β-cyclodextrin (mβCD) hinders the auxin-triggered ROP6 activation. **Bottom panel:** Quantitative analysis of the relative active ROP6 level from three independent experiments. Three-week old transgenic seedlings containing 35S∷GFP-ROP6 were used to prepare protoplast and treated with 0.0001% ethanol (mock), 10mM mβCD for 30 mins, 100nM IAA for 10mins, and 10mM mβCD for 30 mins followed by 10 mins IAA treatment. The MBP-RIC1 bound active ROP6 were detected using anti-GFP antibodies. The relative active ROP6 level was determined as the amount of GTP-bound ROP6 divided by the amount of total GFP-ROP6. The relative ROP activity in different treatments was normalized to that from mock-treated control, and this value was set as standard 1. Data were presented as mean ± SD. **(G,H)** Auxin increases the level of ROP6 S-acylation in a dose-dependent manner. **(G)** Representative Western blot image showing the level of ROP6 S-acylation in response to auxin treatments. S-acylation level of ROP6 proteins was assayed using the acyl-RAC method. For each assay, the sample was compared with the loading control (LC) with or without hydroxylamine (HYD) for hydroxylamine-dependent capture of S-acylated proteins. The samples treated with NaCl instead of HYD was used as the negative control. Three-week-old transgenic plants containing 35S∷GFP-ROP6 were sprayed with different concentrations of IAA (0, 10, 20, 50, 100 nM and 1 μM) and leaf tissues were collected 10 mins after spraying. **(H)** Quantitative analysis of the relative S-acylated ROP6 level from three independent experiments. The relative S-acylated ROP6 level was determined as the amount of S-acylated ROP6 divided by the amount of total GFP-ROP6. The relative S-acylation level in different treatments was normalized to that from mock-treated control as standard 1. Data were presented as mean ± SD.

To understand how ordered lipid domains contribute to the formation of indentation polarity, we examined the effect of sterols on the activation of ROP6, an indentation polarity marker that is preferentially distributed to the indentation region and critical for indentation formation (Fu et al., 2009; Lin et al., 2013; Xu et al., 2010). Previously we showed that ROP6 is activated by auxin (Xu et al., 2014; Xu et al., 2010). We further found that reducing PM sterols by mβCD treatment greatly hindered auxin-induced activation of ROP6 (Fig. 2F). Moreover, auxin increases the amount of *S*-acylated ROP6 in a dose-dependent manner (Figs. 2G and 2H). It was shown that only active but not inactive form of ROP6 was acylated and partitioned into lipid-raft like, detergent-resistant membrane fraction (DRM) (Sorek et al., 2017; Sorek et al., 2010). Taken together, these results suggest auxin promotes the formation of ordered lipid domains required for the indentation polarity establishment.

### Auxin promotes the formation of TMK1 nanoclusters with larger size and slower diffusion

We next investigated how auxin promotes the formation and polarization of ordered lipid domains to the indentation region. Auxin signaling is mediated by at least two major pathways: The nuclear-based TIR1/AFB pathway that regulates the expression of auxin-induced genes (Salehin et al., 2015; Tan et al., 2007) and the PM-based TMK pathway involved in non-transcriptional auxin responses (Cao et al., 2019; Xu et al., 2010). We speculated that the TMK-dependent pathway regulates lipid ordering and its polarity, given its role in the auxin-induced activation of ROP6 in the PM. In addition, the *tmk1tmk4* double mutant showed reduced lipid order in the PM of PCs (Fig. S3). As an initial step in testing our hypothesis, we analyzed the spatiotemporal dynamics of TMK1, a major member of the TMK clade with four functionally overlapping members (Dai et al., 2013; Xu et al., 2014). We used total internal refection fluorescence (TIRF) microscopy and single-particle tracking methods to analyze TMK1 dynamics in 2-3 days-old cotyledon PCs. Indeed, TMK1-GFP, expressed under *TMK1*’s native promoter, was laterally segregated as nano-particles at the PM (Supplementary Video 4 and Fig. 3A). Treatment with 100 nM IAA resulted in a dramatic decrease in the density of TMK1-GFP particles (Fig. 3B) and a significant increase in the particle size (Fig. 3C-3E). These results suggest that auxin promotes the formation of larger TMK1 nanoclusters, possibly by coalescing single proteins or small and unstable nanoclusters. Further diffusion analyses showed that a two-component model describes the distribution of diffusion coefficients much better than a one-component model (Fig. S4), indicating TMK1-GFP particles on the membrane consist of at least two populations, one with slow and one with fast diffusion. IAA treatment (100 nM, 10 min) caused a significant reduction in diffusion coefficient along with an increase in the fraction of the slow-diffusion population of TMK1-GFP particles (Fig.3F, Table. S1). By contrast, the same IAA treatment had no effect on the particle size of GFP-tagged FERONIA, another receptor-like kinase at the PM also known to affect PC morphogenesis (Fig. S5) (Li et al., 2015; Lin et al., 2018). These results suggest that auxin specifically promotes the formation of the TMK nanoclusters with larger size and slower diffusion.

**Fig 3.**
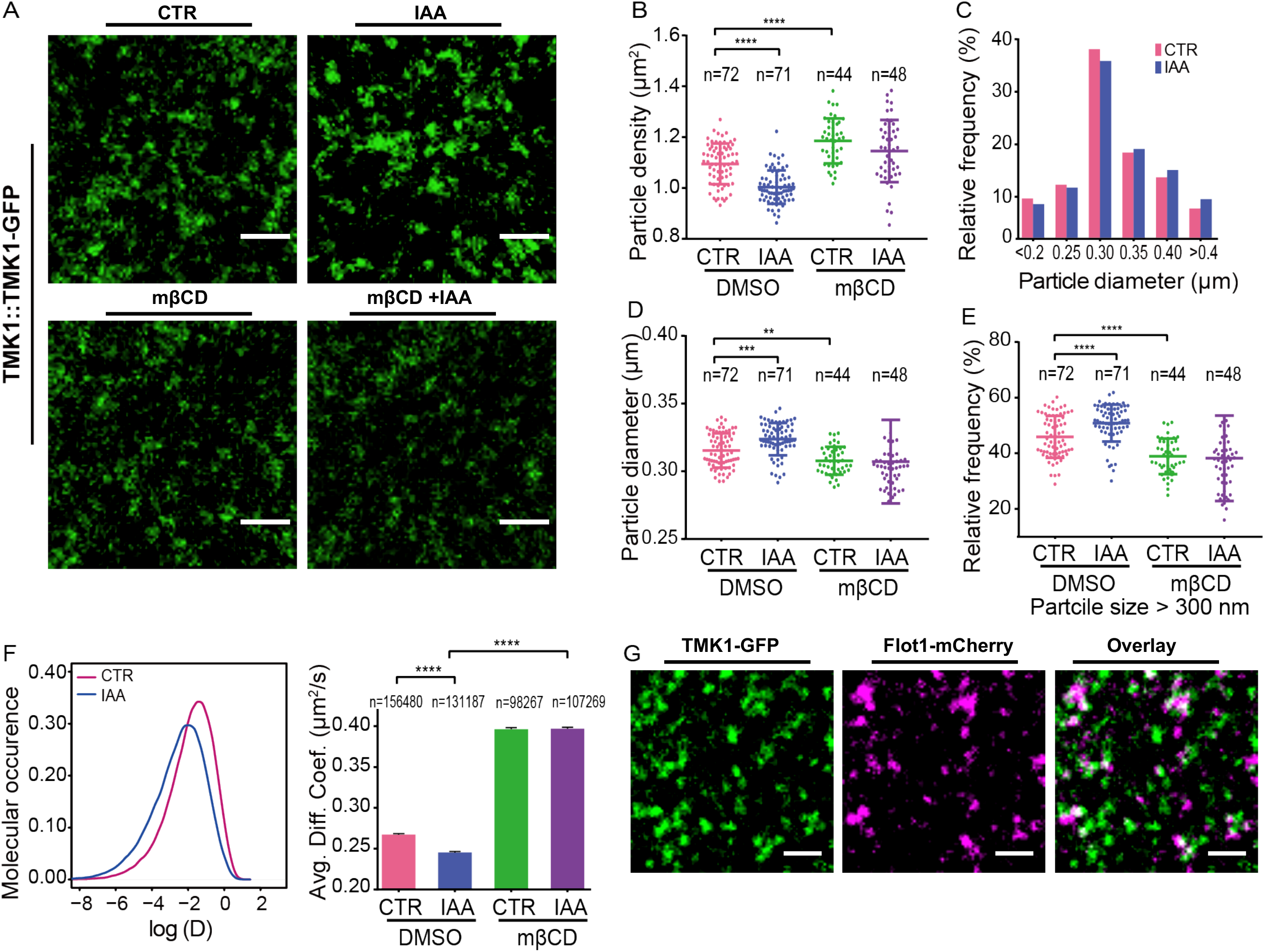
Auxin promotes the TMK1 nanoclustering in a sterol-dependent manner. (**A**) TIRF image of pavement cells fluorescently labeled with TMK1∷TMK1-GFP with or without IAA (100 nM, 10 mins) and methyl-β-cyclodextrin (mβCD, 10mM, 30mins) treatments. Movies were recorded with a 100-ms exposure time for 20s. Over this imaging period photobleaching was negligible. Scale bars = 2 μm **(B)** Quantitative comparison of TMK1 particle densities between control and IAA treatment with or without mβCD pre-treatment. The IAA treatment significantly decreases the number of TMK1 particles at the PM in the absence of mβCD. Data were presented as mean ± SD. *n* represents the number of independent cells. **(C)** Particle size distribution histogram between control and IAA treatments. **(D)** Quantitative comparison of average TMK1 particle size between control and IAA treatment with or without mβCD pretreatment. **(E)** Relative frequency of TMK1 particles with size larger than 300nm between control and IAA treatment with or without mβCD pretreatment. Data were presented as mean ± SD. *n* represents the number of independent cells. **(F, Left panel)** The density curves of diffusion coefficients for TMK1 particles with (blue curve) or without IAA treatment (pink curve). **(F, Right panel)** Comparison of average diffusion coefficients for TMK1 particles with or without IAA and mβCD treatments. Data were presented as mean ± SEM. *n* represents the number of particles. **(G)** Representative TIRF images of pavement cells co-expressing TMK1-GFP and Flot1-mCherry, Scale bars = 2 μm.

### Auxin-triggered TMK1 nanoclustering and lipid ordering are inter-dependent

Because ligand-induced receptor clustering may modulate the local lipid environment and stabilize ordered lipid domains around the receptor (Hofman et al., 2008; Kusumi et al., 2004; Sohn et al., 2006; Stone et al., 2017), we hypothesize that auxin-triggered TMK1 nanoclustering stabilizes a local ordered-lipid environment. To test the hypothesis, we first studied the dynamic relationship between TMK1 and flotillin proteins at the PM. As shown in Fig. 3G, TMK1-GFP particles are at least partially colocalized with Flotillin1-mCherry particles. To functionally test the role of ordered lipid domains in TMK1 nanoclustering, we analyzed TMK1 dynamics in the presence of the sterol-disrupting agent mβCD (10 mM for 30 min). This treatment greatly hindered the auxin-induced increase in sizes (Fig. 3B-E) and reduction in diffusion of TMK1 particles (Fig. 3F and Table. S1). These results indicate that auxin-induced TMK1 nanoclustering are dependent upon sterol content in the PM.

To test whether auxin-induced changes in lipid ordering in the PM requires TMKs, we analyzed the dynamics of flotillin-mVenus under its native promoter, which was used as an indicator for the dynamics of ordered nanodomains at the PM. Similar to TMK1, IAA treatment (100 nM, 10 mins) increased the flotillin-mVenus particle size (Figs. 4A-4D). Diffusion analyses also revealed the presence of a fast-diffusing population and a slow-diffusing population (Fig. S6). Auxin treatment increased the fraction of the slow-diffusing population with a decrease in the fraction of the fast-diffusing population (Figs. 4E and 4F, Table. S2). The diffusion coefficient of flotillin1-mVenus in *tmk1tmk4* double mutant was approximately an order of magnitude faster than that in wild type (Fig. 4F). In addition, the *tmk1/tmk4* double mutant displayed a substantial resistance to the promotion of flotillin particle size and the reduction of particle diffusion triggered by auxin (Figs. 4C and 4D). These data demonstrate that a significant portion of auxin’s effect on the dynamics of flotillin-associated ordered nanodomains is regulated via the TMK1-mediated pathway.

**Fig 4.**
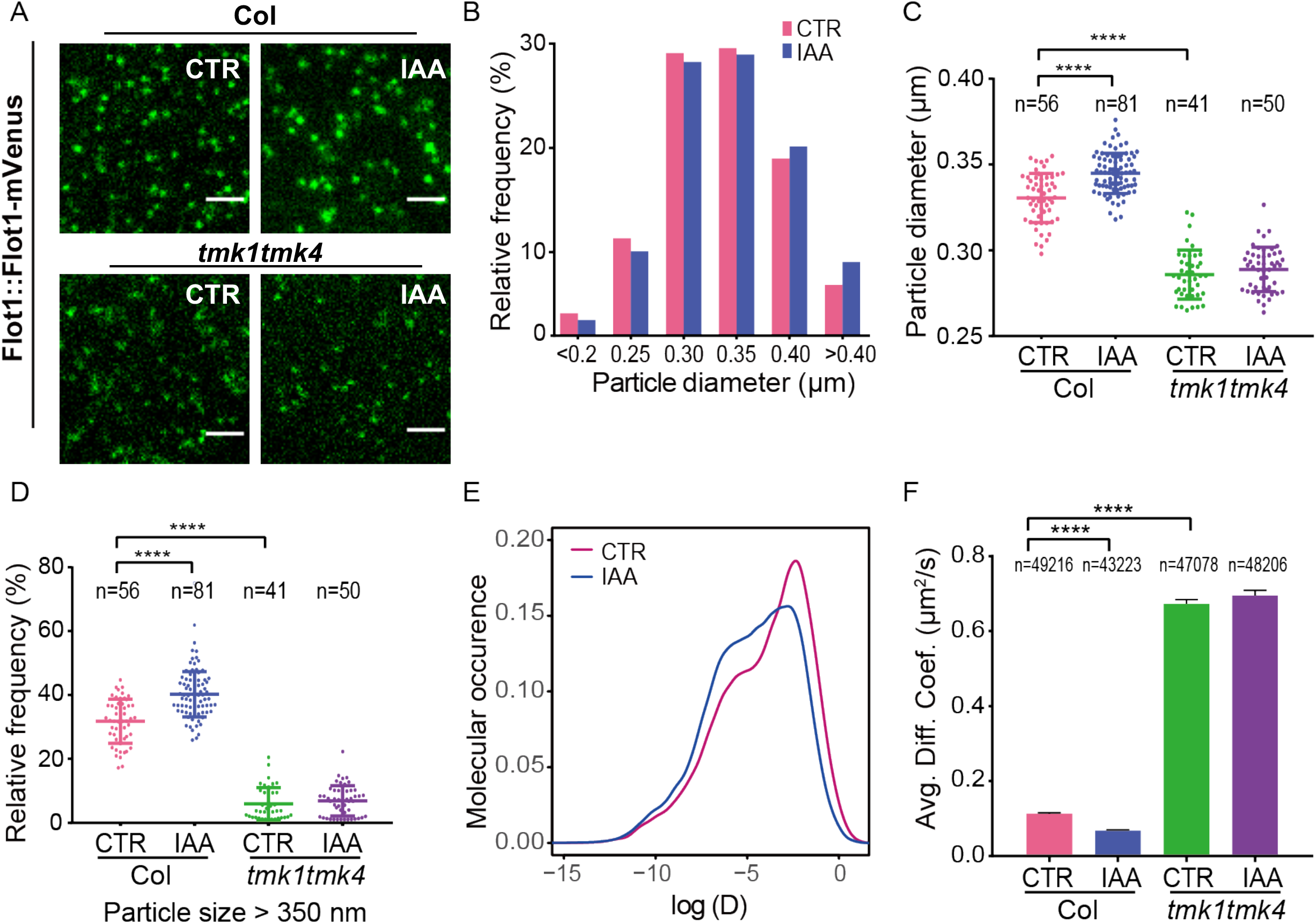
TMK contributes to the auxin-promoted nanoclustering and stability of flotillin particles. **(A)** TIRF images of pavement cells fluorescently labeled with Flot1∷Flot1-mVenus with or without IAA (100 nM, 10 mins) in wild-type *(Col-0)* and *tmk1tmk4* double mutant background. Scale bars = 2 μm. **(B)** Particle size distribution histogram between control and IAA treatment. **(C)** Quantitative comparison of Flot1 particle size between control and IAA treatment in wild-type *(Col-0)* and *tmk1tmk4* double mutant background. **(D)** Relative frequency of Flot1 particles with size larger than 350nm between control and IAA treatment in in wild-type *(Col-0)* and *tmk1tmk4* double mutant background. Data were presented as mean ± SD. *n* represents the number of independent cells. **(E)** The density curves of diffusion coefficients for Flot1 particles with (blue curve) or without IAA treatment (pink curve). **(F)** Comparison of average diffusion coefficients for Flot1 particles without and with 100nM IAA treatment in wild-type *(Col-0)* and *tmk1tmk4* double mutant background. Data were presented as mean ± SEM. *n* represents the number of particles. See methods for detailed statistical analyses.

### TMKs are required for auxin-mediated nanoclustering of ROP6

Our results described above show that auxin promotes TMK1 nanoclustering (Figs. 3A-3E), lipid ordering (Figs. 2A-2D), and the polarization of ordered lipid domains to the indentation sites (Figs. 1A-1E) where ROP6 localization. Moreover, previous studies suggest that ROP6 is activated by auxin in a TMK-dependent manner (Xu et al., 2014; Xu et al., 2010) and we further found that auxin-induced ROP6 activation requires sterols (Fig. 2F). Thus, we hypothesize that auxin promotes the nanoclustering of active ROP6 in a TMK- and sterol-dependent manner. Indeed, TIRF and single-particle tracking analysis showed that the constitutively active (CA) form of ROP6 was preferentially distributed in the particles with larger size and slower-diffusion properties in comparison with the inactive form of ROP6 (Dominant Negative, *DNrop6*, Figs. 5A and 5B, Table. S3). To test the effect of auxin on ROP6 nanoclustering, we performed TIRF and single-particle tracking of EGFP-ROP6 expressed under its native promoter. EGFP-ROP6 showed clear lateral segregation on the PM in the *ROP6p∷EGFP-ROP6* line (Supplementary Video 5, Fig. 5C) and treatment with 100 nM IAA, which induces ROP6 activation by four-fold (Xu et al., 2010), significantly increased particle size (Figs. 5D). Further analyses showed that, like TMK1 particles, the distribution of diffusion coefficients of ROP6 particles can be better described by a two-component model, indicating the existence of at least two populations of ROP6 particles, one slow- and one fast-diffusing population (Fig. S7). Consistently, IAA treatment significantly reduced the ROP6 diffusion coefficient with an increase in the fraction of the slow-diffusion population (Figs. 5E and 5F, Table. S3). Furthermore, the effect of auxin on ROP6 particles was completely abolished in the presence of sterol-depleting agent mβCD (Figs. 5G and 5H). These data suggest that auxin-triggered ROP6 activation induces ROP6 stabilization in sterol-dependent PM nanodomains with increased size and reduced diffusion.

**Fig 5.**
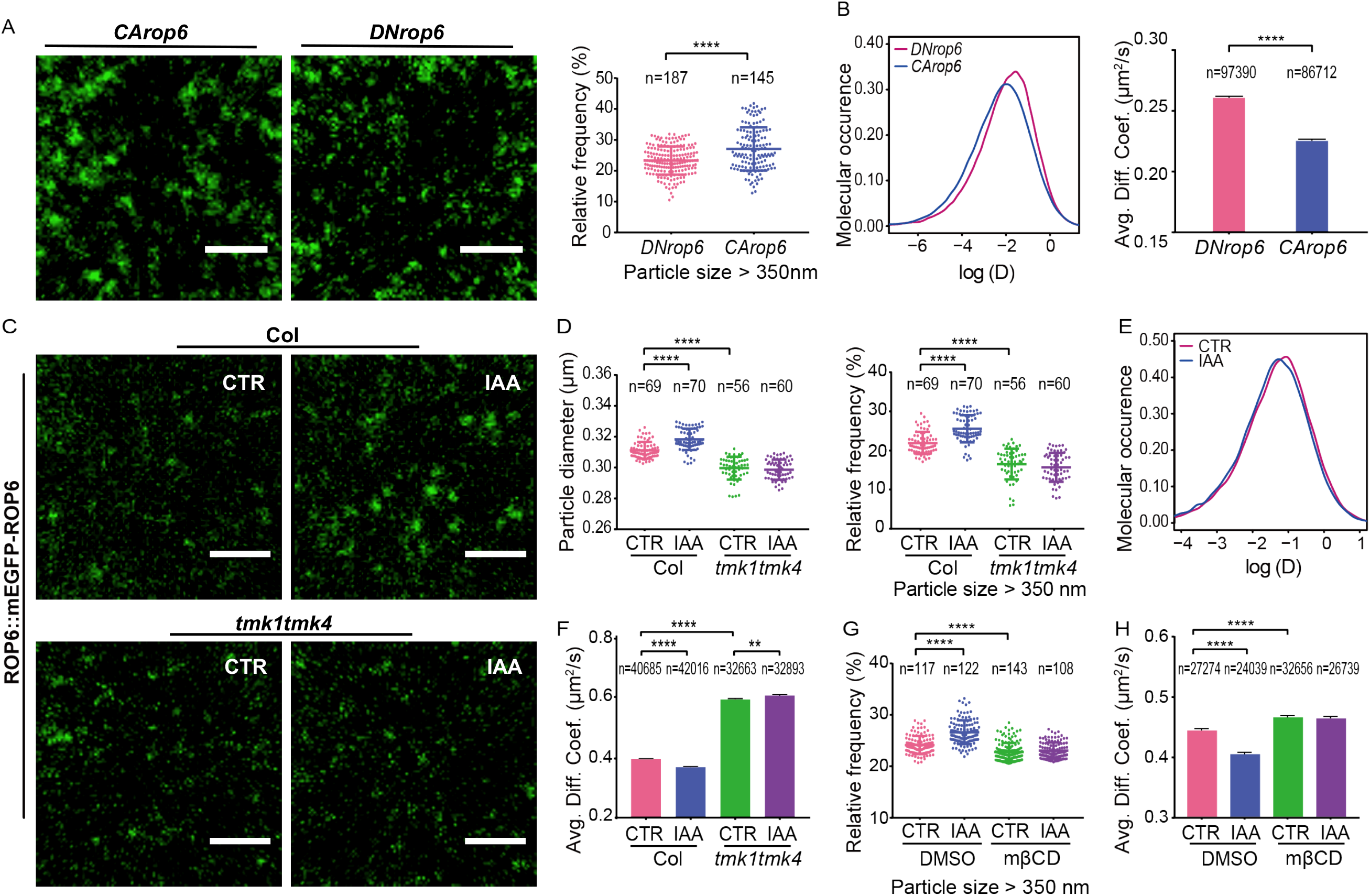
TMK contributes to the auxin-promoted nanoclustering and stability of ROP6 particles. **(A, B)** *CArop6* forms larger particles with slower diffusion rate than *DNrop6* at the plasma membrane. **(A, left panel)** Representative TIRF images of pavement cells fluorescently labeled with 35S∷GFP-*CArop6* and 35S∷GFP-*DNrop6*. Scale bars = 2μm. **(A, right panel)** Relative frequency of particles with size larger than 350nm between GFP-*CArop6* and GFP-*DNrop6*. *n* represents the number of independent cells. Data were presented as mean ± SD. **(B) Left panel:** The density curves of diffusion coefficients for GFP-*CArop6* (blue curve) and GFP-*DNrop6* (pink curve) particles. **Right panel:** Comparison of average diffusion coefficients for GFP-*CArop6* and GFP-*DNrop6* particles. Data were presented as mean ± SEM. *n* represents the number of particles. **(C-F)** Auxin-induced increase in size and decrease in diffusion rate of ROP6 particles is compromised in the *tmk1tmk4* double mutant. (**C**) TIRF image of pavement cells fluorescently labeled with ROP6∷GFP-ROP6 with or without IAA (100 nM, 10 mins) in wild-type *(Col-0)* and *tmk1tmk4* double mutant background. Scale bars = 2 μm. **(D) Left panel:** Quantitative comparison of average ROP6 particle size between control and IAA treatment in wild-type *(Col-0)* and *tmk1tmk4* double mutant background. **Right panel:** Relative frequency of ROP6 particles with size larger than 350nm between control and IAA treatment in wild-type *(Col-0)* and *tmk1tmk4* double mutant background. Data were presented as mean ± SD. *n* represents the number of independent cells. **(E)** The density curves of diffusion coefficients for ROP6 particles with (blue curve) or without IAA treatment (pink curve). **(F)** Comparison of average diffusion coefficients for ROP6 particles without and with 100nM IAA treatment in wild-type *(Col-0)* and *tmk1tmk4* double mutant background. Data were presented as mean ± SEM. *n* represents the number of particles. See methods for detailed statistical analyses. **(G, H)** Auxin-induced increase in size and decrease in diffusion rate of ROP6 particles is abolished in the presence of the sterol-depleting agent methyl-β-cyclodextrin (mβCD). **(G)** Relative frequency of ROP6 particles with size larger than 350nm between control and IAA treatment with or without mβCD pretreatment. Data were presented as mean ± SD. *n* represents the number of independent cells. **(H)** Comparison of average diffusion coefficients for ROP6 particles with or without IAA and mβCD treatments. Data were presented as mean ± SEM. *n* represents the number of particles.

Given the critical role of TMK1/4 in the auxin-induced stabilization of ordered lipid nanodomains (Fig. 4), we suspected that TMK1/4 also regulates behavior of ROP6 nanoclusters. Indeed, we found that in the *tmk1tmk4* double mutants the ROP6 particles are much smaller in size (Fig. 5D) and have a higher mobility (Figs. 5E and 5F). In addition, the auxin-triggered particle size increase (Fig. 5D) and diffusion decrease (Figs. 5E and 5F) were greatly compromised in the *tmk1tmk4* double mutant, indicating that TMK1/4 are critical for auxin-induced changes in the dynamic behavior of ROP6 particles. Taken together, these data strongly support the hypothesis that TMK1 regulates auxin-induced ROP6 nanoclustering via shaping the sterol-rich lipid environment at the PM. Interestingly, the auxin-induced ROP6 nanoclustering has also been described in Arabidopsis roots by Jaillais’s group, who showed that phosphatidylserine lipid is important for auxin-mediated ROP6 nanoclustering and phosphatidylserine biosynthesis mutants exhibit severe PC morphogenesis defect (Platre et al., 2019). Thus, ROP6 nanoclusters in PCs may be enriched in both sterols and phosphatidylserines.

### Modeling predicts a crucial role for a CMT-based nanoclustering feedback loop in polarity establishment

Our experimental data (above) show that auxin induces both indentation polarity and sterol-dependent nanoclustering of TMK1 and ROP6 and that sterol-based lipid ordering is critical for the establishment of indentation polarity. These results beg the question: How does auxin-induced nanoclustering of auxin signaling components contribute to the polarity establishment during PC morphogenesis? To address this question, we constructed a mathematical model describing the process of early polarity establishment in PCs. We first considered the network including TMK1 nanoclusters, ordered lipid nanodomains, and ROP6 nanoclusters on a closed moving interface representing PM in a two-dimensional space (cross-section). TMK1 nanoclusters, ordered lipid domains, and ROP6 nanoclusters are all modeled as diffusible molecules on the PM by using reaction-diffusion equations. The moving interface is computed by using level set method (Osher, 1988) evolved under a specific velocity field. Our initial model (Fig. 6A) incorporates the assumptions based on the experimental observations that: (i) TMK1 nanoclusters and flotillin-associated ordered lipid nanodomains restrict the diffusion of each other, (ii) TMK1-stablized ordered lipid nanodomains promote ROP6 nanoclustering and activation, (iii) ordered lipid nanodomains restrict the diffusion of ROP6, (iv) the cell is expanding in an isotropic way due to some additional signals, which are not included as variables in the model. As shown in Fig. 6A, nanoclustering of TMK1 and ROP6 proteins alone is insufficient to generate polarity, suggesting that other events or processes are also required for PC polarity establishment.

**Fig 6.**
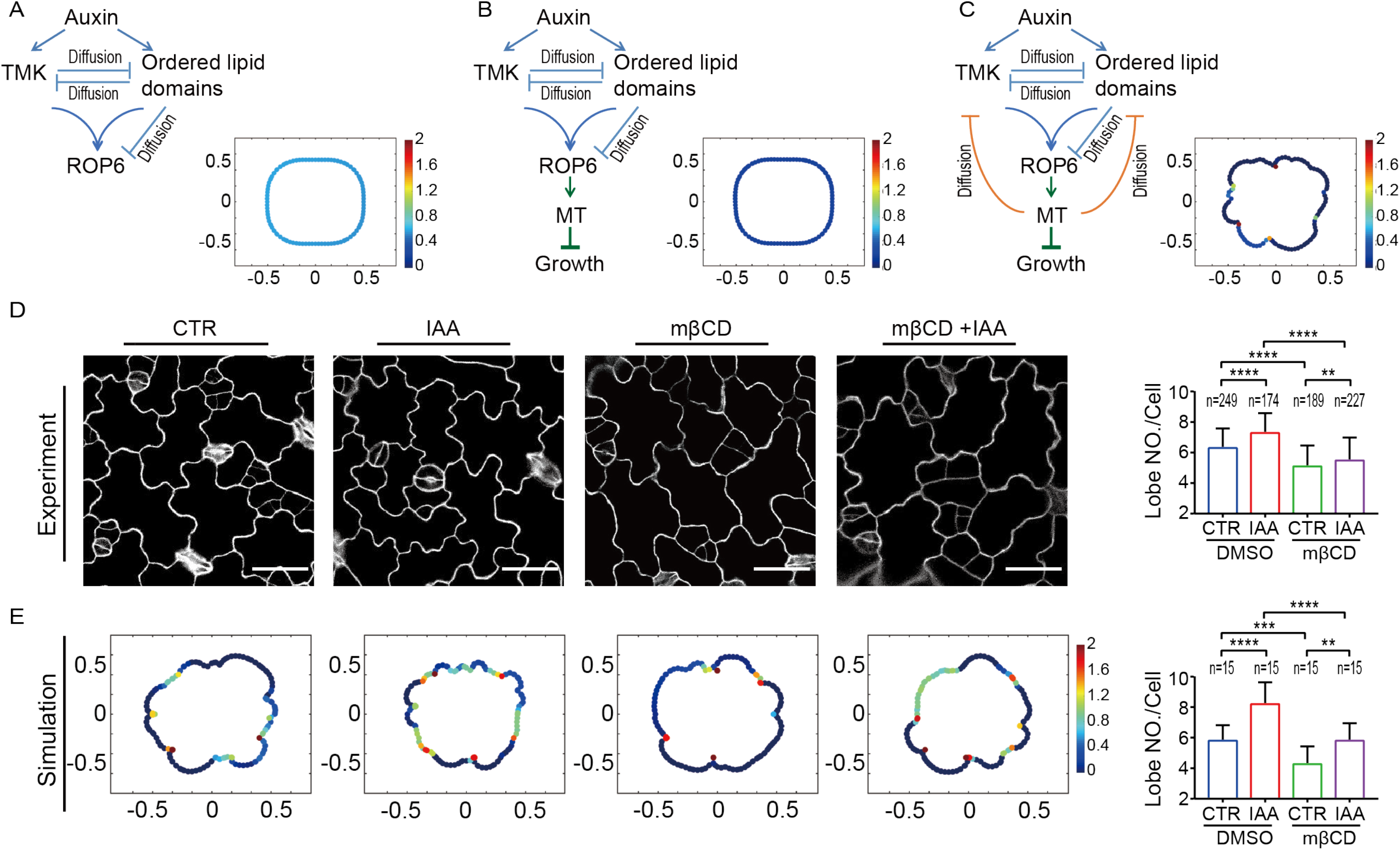
A model of microtubule-based nanoclustering for pavement cell polarity establishment. **(A) Left Panel:** Schematic diagram for the computational model without the involvement of cortical microtubules (CMTs). **Right Panel:** Simulated image of the pavement cell generated from the model shown on the left. The color scale indicates the level of active ROP6. **(B) Left Panel:** Schematic diagram for the computational model with the addition of CMT-mediated growth restriction. **Right Panel:** Simulated image of the pavement cell generated from the model shown on the left. **(C) Left panel:** Schematic diagram for the computational model with the CMT-based nanoclustering feedback loop. **Right panel:** Simulated image of the pavement cell generated from the model shown on the left. Also see Supplementary Video 6. **(D, E)** Experimental and simulation data consistently show that auxin-mediated pavement cell interdigitation is compromised in the presence of 10mM methyl-β-cyclodextrin (mβCD). **(D) Left panels:** Representative confocal images of pavement cells in wild-type (Col-0) cotyledons grown on solid ½ MS medium supplemented with DMSO (mock treatment; 1^st^ panel), 50 nM IAA (2^nd^ panel), 10mM mβCD (3^rd^ panel) and 50nM IAA plus 10mM mβCD (4^th^ panel). Scale bars = 15 μm. **Right panel:** Quantitative analysis of the number of lobes under different treatments. *n* represents the number of independent cells. **(E) Left panels:** Representative images of simulated pavement cells with altered levels of auxin and sterol-enriched nanodomains. The color scale indicates the level of active ROP6. More simulated images are presented in Figs. S10-S13. **Right panel:** Quantitative analysis of the number of lobes under different simulations. *n* represents the number of simulated cells. Data were presented as mean ± SD. See methods for detailed statistical analyses.

ROP6 signaling promotes cortical microtubule (CMT) organization (Fu et al., 2005; Fu et al., 2009) during PC development, and CMT then impacts cell shape by directing the deposition of cellulose microfibrils, which resist cell expansion resulting from internal turgor pressure (Bringmann et al., 2012; Hamant and Traas, 2010). Hence, we tested the model (Fig. 6B) with CMT slowing isotropic cell expansion by introducing CMT as an additional network component recruited by ROP6. As shown in Fig. 6B, addition of CMT-mediated growth restriction into the model still fails to generate polarity, indicating that a key component in our model is missing. We next considered a potential role for CMTs in limiting the diffusion of TMK1 nanoclusters and ordered lipid nanodomains as a feedback loop in addition to slowing down the cell expansion (Fig. 6C). CMTs are physically attached to the inner leaflet of the PM, and thus could presumably hinder the mobility of TMK1 nanoclusters and ordered lipid nanodomains. Interestingly, we found that constitutively active ROP6 was localized along CMTs, which was confirmed by the oryzalin treatment (Fig. S8), suggesting a potential physical connection between CMTs and active ROP6s. Thus, we incorporated into our model the assumption that CMTs restrict the diffusion of TMK1 nanoclusters and ordered lipid nanodomains as a form of feedback regulation (Fig. 6C). Using this model (Fig. 6C), the cell underwent the morphological change from the initial condition of a rounded or rectangular shape, to a shape with multi-polarity, as we observe in PCs (Supplementary Video 6). Hence our modeling predicts that both TMK1-based nanodomain formation and its stabilization by the CMT-based feedback loop are necessary and sufficient for the generation of multi-polarity in PCs.

We further incorporated auxin effect into the model by proportionally increasing the initial concentration of TMK1 nanoclusters and ordered lipid nanodomains, where the fold change between control and auxin treatment is set according to our experimental data (see “Materials and Methods” section). Our auxin-triggered nanocluster-based polarity model (Fig 6C) was validated by existing and new experimental data. First, our model successfully reproduced the dose-dependent effect of auxin on PC interdigitation as previously described (Xu et al., 2010) (Fig. S9). Our model explains why PC interdigitation requires an optimal level of auxin (Xu et al., 2010). The modeling shows that the number of polarity sites increases with increasing auxin concentration up to an optimum level. However, when the auxin level is higher than the optimal concentration, a very high number of auxin-induced TMK1/sterol-rich nanodomains starts to coalesce and saturate the PM, resulting in a low degree of lateral segregation of ROP6 nanodomains, and eventually less polarity sites (Supplementary Video 7). Second, the role of ordered lipid nanodomains in polarity establishment was confirmed using our model. In the simulations, a decrease in the initial concentration of ordered lipid nanodomains resulted in the reduced number of polarization sites (as indicated by the number of lobes) (Fig. 6E and Figs. S10-S13). Consistently, our experimental findings show that genetic (Fig1F and Fig. S2) and chemical perturbations (Fig. 6D) of sterol biosynthesis significantly reduced the PC interdigitation. Lastly, our model predicts that auxin-promoted PC interdigitation is greatly compromised when the level of ordered nanodomains is reduced (Fig. 6E and Figs. S10-S13). These computational results are consistent with our experimental observations as shown in Fig. 6D.

### CMTs stabilize sterol-dependent TMK1 and ordered lipid nanodomains

An important prediction from our model is that CMTs stabilize TMK1 nanoclusters and ordered lipid nanodomains. To observe an immediate response of microtubule disruption, we treated the cotyledons expressing TMK1 or flotillin 1 with high concentration of oryzalin (5mM) for 30 mins. As shown in Fig. 7A-7C, microtubule disruption with oryzalin greatly decreased the size and increased the diffusion of TMK1 particles on the PC. Flotillin particles became highly diffusive upon oryzalin treatment, even though no significant changes in particle size were observed (Fig. 7D-7F). In addition, depolymerizing CTMs with oryzalin did not block the auxin-induced diffusion decrease of TMK1 and flotillin particles, but led to smaller particles with faster diffusion compared to the control condition (with intact CTMs) upon auxin treatment (Fig. 7A-7F), supporting that CTMs act downstream of auxin-induced initial nanoclustering to further stabilize TMK1 and flotillin clusters. Furthermore, our computational model is unable to produce cell polarity unless the CMT-dependent feedback regulation on nanocluster diffusion restriction was present (Figs. 6A-C). This suggests that CMT-dependent nanodomain stabilization is essential for polarity establishment.

**Fig 7.**
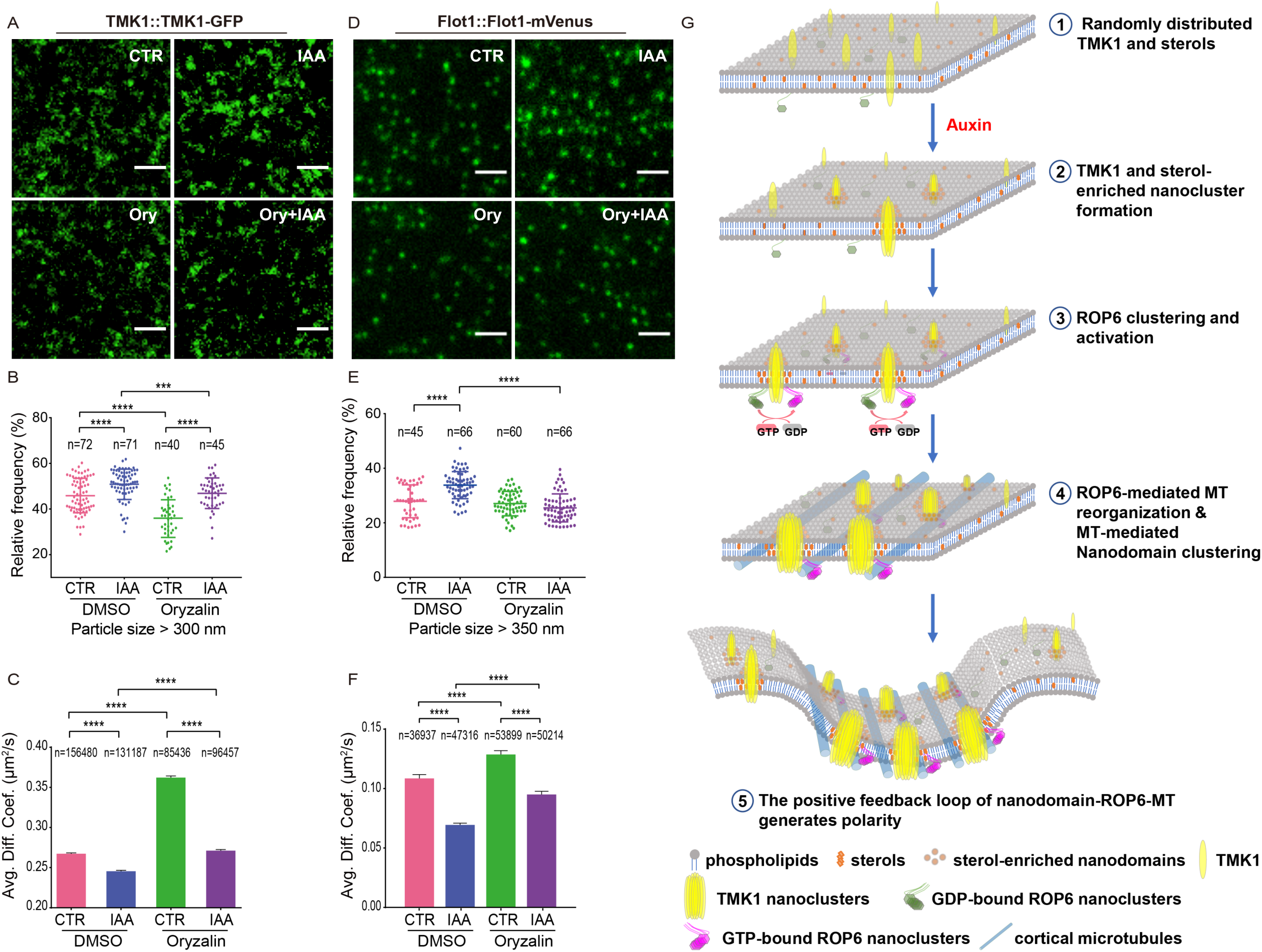
Microtubule-involved promotion of TMK1 and flotillin nanoclustering at the PM is essential for multi-polarity establishment. **(A-C)** Auxin effect on dynamic change of TMK1 particles is compromised by microtubule disruption. **(A)** TIRF image of pavement cells fluorescently labeled with TMK1∷TMK1-GFP with or without IAA (100 nM, 10 mins) and oryzalin (5μM, 30mins) treatments. Scale bars = 2 μm. **(B)** Relative frequency of TMK1 particles with size larger than 300nm between control and IAA treatment with or without oryzalin pre-treatment. Data were presented as mean ± SD. *n* represents the number of independent cells. **(C)** Comparison of average diffusion coefficients for TMK1 particles between control and IAA treatment with or without oryzalin pre-treatment. Data were presented as mean ± SE. *n* represents the number of particles. Note: The oryzalin and mβCD treatments (Fig. 3) on TMK1 dynamics were performed in the same batch and the same mock treatment was used. The data and representative images for the control group are the same for Fig. 3 and Figs. 7A-7C. **(D-F)** Auxin effect on dynamic change of flotillin particles is compromised by microtubule disruption. **(D)** TIRF image of pavement cells fluorescently labeled with Flot1∷Flot1-mVenus with or without IAA (100 nM, 10 mins) and oryzalin (5μM, 30mins) treatments. Scale bars = 2 μm **(E)** Relative frequency of Flot particles with size larger than 350nm between control and IAA treatment with or without oryzalin pre-treatment. Data were presented as mean ± SD. *n* represents the number of independent cells. **(F)** Comparison of average diffusion coefficients for Flot1 particles between control and IAA treatment with or without oryzalin pre-treatment. Data were presented as mean ± SE. *n* represents the number of particles. See methods for detailed statistical analyses. **(G)** Schematic illustration showing the steps involved in the auxin-mediated multi-polarity establishment. TMK proteins at the PM are first induced by auxin to form large nanoclusters, which is accompanied by the formation of sterol-rich ordered lipid nanodomains. The newly formed large lipid nanodomains provides an ideal lipid environment for ROP6 nanoclustering and activation. The active ROP6 interacts with downstream effectors and promotes the cortical microtubule (CMT) ordering, which further restrict the diffusion of TMK1 nanoclusters and their surrounding ordered lipid nanodomains. The positive feedback loop of ordered lipid nanodomain-ROP6 signaling-CMT ordering eventually leads to the establishment of indentation polarity. Proteins and structures are not to scale.

Taken together, we propose the following model of multi-polarity establishment (Fig. 7G) in which: 1) Upon auxin signaling, the order lipid nanodomain-preferring TMK1 proteins are clustered. TMK1 nanoclustering further promotes the formation of larger, stabilized ordered lipid nanodomains, probably by coalescing small, unstable lipid nanodomains associated with TMK1 proteins. 2) The newly formed large lipid nanodomains provides an ideal lipid environment for ROP6 nanoclustering and activation. 3) The active ROP6 interacts with downstream effectors and promotes CMT ordering, which further restrict the diffusion of TMK1 nanoclusters and their surrounding ordered lipid nanodomains. 4) The initial asymmetric distribution of active ROP6 is further amplified by the positive feedback loop of ordered lipid nanodomain-ROP6 signaling-CMT ordering, and eventually leads to a stable multi-polarity establishment.

## Conclusion and Discussion

Our findings reveal a nanodomain-centric principle for the establishment of cell polarity and for auxin signaling at the plasma membrane. These findings demonstrate a role for nanoclustering in controlling cell polarity formation, expanding its roles in the regulation of fundamental processes such as cell spreading (Kalappurakkal et al., 2019). Importantly our findings have established a new paradigm for cell polarity establishment that can explain polarization that is self-organizing or induced by uniform or localized signals. Local protein recruitment and/or endomembrane trafficking have been prevailing models to explain cell polarization (Butler and Wallingford, 2017; Nelson, 2003; Strutt and Strutt, 2009; Vladar et al., 2009). Here we demonstrate that the coordination of nanoclustering of cell surface signaling components with microtubule-dependent feedback regulation of nanocluster diffusion restriction is necessary and sufficient for the establishment of cell polarity induced by a uniform field of auxin. This coordinated action of ligand-induced nanoclustering and microtubule-imposed diffusion restriction likely provides a common design principle underlying polarity establishment in different cell systems.

Many cell polarity proteins such as Rac1 and Cdc42 are found in nanoclusters (Maxwell et al., 2018; Meca et al., 2019; Remorino et al., 2017; Sartorel et al., 2018), but it is unclear how their nanoclustering modulates cell polarity establishment. Here we integrate data-based modeling with model-inspired experiments to show that auxin-triggered nanoclustering of TMKs and ROP6 is required but insufficient for the establishment of cell polarity in PCs, whereas the coordination of nanoclustering with diffusion restriction of nanoclusters by auxin-triggered CMT ordering enables PCs to establish the multi-polarity required for their characteristic shape. ROP6 promotes the organization of ordered CMTs, which are attached to the PM via anchor proteins. Our modeling predicts and experiments verify that CMTs slow the diffusive movement of TMK1 and flotillin particles at the PM, which might enhance their collision and clustering. Such microtubule-dependent feedback regulation of nanoclusters is critical for the generations of PC polarity (Fig. 7). This feedback loop-dependent regulation of nanocluster is distinct from the TMK-based ROP6 nanoclustering directly triggered by auxin but is essential for the generation of PC polarity. The CMT regulation of ROP6 nanoclustering appears to be mediated by the direct association of active ROP6 (in the nanoclusters) with CMTs, as we found active ROP6 is associated with CMTs as nanoclusters. A critical role for microtubules in cell polarization is well recognized in eukaryotic cells (Drubin and Nelson, 1996; Siegrist and Doe, 2007; St Johnston, 2018), and their association with Rho GTPase signaling proteins (ICR1, MDD1, RhoGEFs, etc) is widely documented (Fu et al., 2009; Hoefle et al., 2011; Oda and Fukuda, 2012; Ren et al., 1998). CMTs have been implicated in regulating PIN protein polarization and ROP6 nanoclustering apparently also impacts PIN polarity (Heisler et al., 2010; Kleine-Vehn et al., 2008; Platre et al., 2019). During symmetry breaking at starvation exit in fission yeast cells, microtubules are important for the formation of polarized sterol-rich membrane domains (Makushok et al., 2016). Previous studies have also implicated CMTs in restricting the diffusion of PM-associated signaling molecules, thus, promoting their clustering during signaling (Jaqaman et al., 2011; Lv et al., 2017). All these observations potentially link cytoskeletal modulation of signaling protein nanoclusters to cell polarity formation in various systems. Therefore, future studies should determine whether the coordination of ligand-activated nanoclustering and cytoskeleton-based diffusion restriction of nanoclusters serves as a general principle behind the establishment of cell polarity.

Emerging evidence suggests that ligand-induced formation of protein-lipid nanodomains at the plasma membrane serves as pivotal signaling platforms throughout eukaryotic cells. Here we show that auxin-induced formation of TMK1 nanoclusters and ordered-lipid nanodomains are interdependent. We propose that this ligand-mediated self-organizing lipid-protein interaction provides an important mechanism to generate a nanodomain signaling platform. This is in line with the ligand-induced transmembrane receptor sculpting of the surrounding ordered-lipid environment proposed for animal cells (Hofman et al., 2008; Kusumi et al., 2004; Sohn et al., 2006; Stone et al., 2017). Such clustering of ordered-lipid-associated proteins modulates the activity of signaling proteins associated with the inner leaflet of the PM such as ROP6 GTPase as we showed here and other GTPases (Eisenberg et al., 2011; Eisenberg et al., 2006) in animal systems. Prenylation of these GTPases is a prerequisite for their membrane association, but this association is weak and likely is involved in the membrane association of inactive ROP6 in a much less stable cluster (Figs. 5A and 5B). We showed that auxin promotes ROP6 S-acylation, and it was found that only active but not inactive ROP6 is acylated (Sorek et al., 2017; Sorek et al., 2010). Thus, we propose that the auxin-activated prenylated ROP6 is further modified by acyl (e.g., palmitoyl) groups, which then interact with auxin-induced sterol nanodomains to form more stable ROP6 nanoclusters. Notably, auxin-induced ROP6 nanoclustering also requires phosphatidylserine lipid (negatively charged) and positively charged residues located at the C-terminus of ROP6 in root cells (Platre et al., 2019). Because phosphatidylserine synthesis defects alter PC shape, it is likely phosphatidylserine-dependent ROP6 nanocluster provides another layer of modulation to promote ROP6 nanoclustering. It is worth noting that, in mammalian cells, Ras nanoclustering is regulated by the ratio of cholesterols and phosphatidylserine on the PM in mammalian cells (Zhou and Hancock, 2015; Zhou et al., 2014). In addition, phosphatidylserine coalesces with cholesterols to support phase separation in the model membrane (Maekawa and Fairn, 2015) and is essential for the generation and stabilization of cholesterol-dependent nanoclusters in live cell membrane (Raghupathy et al., 2015). Thus, future studies should elucidate how charged lipid environment coordinates with the sterol-based lipid environment for ligand-mediated GTPase nanoclustering in various systems.

Our findings here and those reported by Jaillais’s group (Platre et al., 2019) together show that auxin-induced nanoclustering of cell surface signaling components plays an important role in the regulation of auxin responses. This PM-localized auxin signaling system enables rapid regulation of cytoplasmic activities and complements the TIR1/AFB-based signaling that mainly controls nuclear gene expression (Salehin et al., 2015; Tan et al., 2007), thus, greatly expanding the toolbox for auxin signaling machinery that is needed for the wide range of auxin responses. Our results show that auxin-induced ROP6 acylation and TMK1-modulated lipid ordering are critical for the nanoclustering of the cell surface auxin signaling components. These findings and the complementary work by Jaillais’s group showing that auxin induces phosphatidylserine-mediated ROP6 nanoclustering in Arabidopsis roots (Platre et al., 2019); altogether, have firmly established a new cell surface auxin signaling mechanism that depends on a localized lipid-protein interaction platform as signaling “hot spot”. Interestingly, auxin also promotes the cleavage of intracellular kinase domain from the cell surface TMKs to regulate nuclear gene expression (Cao et al., 2019). Therefore, the PM-originated signaling mediated by TMKs is emerging as an important auxin signaling mechanism, and thus it would be interesting to determine whether all TMK-dependent auxin signaling pathways are mediated via nanoclustering.

## Acknowledgements

We thank members of the Yang laboratory for their technical assistance and helpful discussions. This work is in part funded by the U.S. National Institute of General Medical Sciences to Z.Y. (GM081451). X.P. was a Natural Sciences and Engineering Research Council of Canada (NSERC) post-doctoral fellowship holder in 2015-2017.

## References

Ariotti, N., Liang, H., Xu, Y., Zhang, Y., Yonekubo, Y., Inder, K., Du, G., Parton, R.G., Hancock, J.F., and Plowman, S.J. (2010). Epidermal growth factor receptor activation remodels the plasma membrane lipid environment to induce nanocluster formation. Molecular and cellular biology 30, 3795–3804.

Bringmann, M., Li, E., Sampathkumar, A., Kocabek, T., Hauser, M.T., and Persson, S. (2012). POM-POM2/cellulose synthase interacting1 is essential for the functional association of cellulose synthase and microtubules in Arabidopsis. The Plant cell 24, 163–177.

Bucherl, C.A., Jarsch, I.K., Schudoma, C., Segonzac, C., Mbengue, M., Robatzek, S., MacLean, D., Ott, T., and Zipfel, C. (2017). Plant immune and growth receptors share common signalling components but localise to distinct plasma membrane nanodomains. eLife 6.

Butler, M.T., and Wallingford, J.B. (2017). Planar cell polarity in development and disease. Nature reviews Molecular cell biology 18, 375–388.

Cao, M., Chen, R., Li, P., Yu, Y., Zheng, R., Ge, D., Zheng, W., Wang, X., Gu, Y., Gelova, Z., et al. (2019). TMK1-mediated auxin signalling regulates differential growth of the apical hook. Nature 568, 240–243.

Chapman, E.J., and Estelle, M. (2009). Mechanism of auxin-regulated gene expression in plants. Annual review of genetics 43, 265–285.

Chen, J., Wang, F., Zheng, S., Xu, T., and Yang, Z. (2015). Pavement cells: a model system for non-transcriptional auxin signalling and crosstalks. Journal of experimental botany 66, 4957–4970.

Dai, N., Wang, W., Patterson, S.E., and Bleecker, A.B. (2013). The TMK subfamily of receptor-like kinases in Arabidopsis display an essential role in growth and a reduced sensitivity to auxin. PloS one 8, e60990.

Dindas, J., Scherzer, S., Roelfsema, M.R.G., von Meyer, K., Muller, H.M., Al-Rasheid, K.A.S., Palme, K., Dietrich, P., Becker, D., Bennett, M.J., et al. (2018). AUX1-mediated root hair auxin influx governs SCF(TIR1/AFB)-type Ca(2+) signaling. Nature communications 9, 1174.

Dinic, J., Riehl, A., Adler, J., and Parmryd, I. (2015). The T cell receptor resides in ordered plasma membrane nanodomains that aggregate upon patching of the receptor. Scientific reports 5, 10082.

Drubin, D.G., and Nelson, W.J. (1996). Origins of cell polarity. Cell 84, 335–344.

Eisenberg, S., Beckett, A.J., Prior, I.A., Dekker, F.J., Hedberg, C., Waldmann, H., Ehrlich, M., and Henis, Y.I. (2011). Raft protein clustering alters N-Ras membrane interactions and activation pattern. Molecular and cellular biology 31, 3938–3952.

Eisenberg, S., Shvartsman, D.E., Ehrlich, M., and Henis, Y.I. (2006). Clustering of raft-associated proteins in the external membrane leaflet modulates internal leaflet H-ras diffusion and signaling. Molecular and cellular biology 26, 7190–7200.

Etienne-Manneville, S., and Hall, A. (2002). Rho GTPases in cell biology. Nature 420, 629–635.

Fendrych, M., Akhmanova, M., Merrin, J., Glanc, M., Hagihara, S., Takahashi, K., Uchida, N., Torii, K.U., and Friml, J. (2018). Rapid and reversible root growth inhibition by TIR1 auxin signalling. Nature plants 4, 453–459.

Fu, Y., Gu, Y., Zheng, Z., Wasteneys, G., and Yang, Z. (2005). Arabidopsis interdigitating cell growth requires two antagonistic pathways with opposing action on cell morphogenesis. Cell 120, 687–700.

Fu, Y., Xu, T., Zhu, L., Wen, M., and Yang, Z. (2009). A ROP GTPase signaling pathway controls cortical microtubule ordering and cell expansion in Arabidopsis. Current biology : CB 19, 1827–1832.

Garcia-Parajo, M.F., Cambi, A., Torreno-Pina, J.A., Thompson, N., and Jacobson, K. (2014). Nanoclustering as a dominant feature of plasma membrane organization. Journal of cell science 127, 4995–5005.

Gerbeau-Pissot, P., Der, C., Grebe, M., and Stanislas, T. (2016). Ratiometric Fluorescence Live Imaging Analysis of Membrane Lipid Order in Arabidopsis Mitotic Cells Using a Lipid Order-Sensitive Probe. Methods in molecular biology 1370, 227–239.

Glebov, O.O., Bright, N.A., and Nichols, B.J. (2006). Flotillin-1 defines a clathrin-independent endocytic pathway in mammalian cells. Nature cell biology 8, 46–54.

Gomez-Mouton, C., Abad, J.L., Mira, E., Lacalle, R.A., Gallardo, E., Jimenez-Baranda, S., Illa, I., Bernad, A., Manes, S., and Martinez, A.C. (2001). Segregation of leading-edge and uropod components into specific lipid rafts during T cell polarization. Proceedings of the National Academy of Sciences of the United States of America 98, 9642–9647.

Hamant, O., and Traas, J. (2010). The mechanics behind plant development. The New phytologist 185, 369–385.

Harding, A.S., and Hancock, J.F. (2008). Using plasma membrane nanoclusters to build better signaling circuits. Trends in cell biology 18, 364–371.

Heisler, M.G., Hamant, O., Krupinski, P., Uyttewaal, M., Ohno, C., Jonsson, H., Traas, J., and Meyerowitz, E.M. (2010). Alignment between PIN1 polarity and microtubule orientation in the shoot apical meristem reveals a tight coupling between morphogenesis and auxin transport. PLoS biology 8, e1000516.

Hoefle, C., Huesmann, C., Schultheiss, H., Bornke, F., Hensel, G., Kumlehn, J., and Huckelhoven, R. (2011). A barley ROP GTPase ACTIVATING PROTEIN associates with microtubules and regulates entry of the barley powdery mildew fungus into leaf epidermal cells. The Plant cell 23, 2422–2439.

Hofman, E.G., Ruonala, M.O., Bader, A.N., van den Heuvel, D., Voortman, J., Roovers, R.C., Verkleij, A.J., Gerritsen, H.C., and van Bergen En Henegouwen, P.M. (2008). EGF induces coalescence of different lipid rafts. Journal of cell science 121, 2519–2528.

Hutten, S.J., Hamers, D.S., Aan den Toorn, M., van Esse, W., Nolles, A., Bucherl, C.A., de Vries, S.C., Hohlbein, J., and Borst, J.W. (2017). Visualization of BRI1 and SERK3/BAK1 Nanoclusters in Arabidopsis Roots. PloS one 12, e0169905.

Jang, J.C., Fujioka, S., Tasaka, M., Seto, H., Takatsuto, S., Ishii, A., Aida, M., Yoshida, S., and Sheen, J. (2000). A critical role of sterols in embryonic patterning and meristem programming revealed by the fackel mutants of Arabidopsis thaliana. Genes & development 14, 1485–1497.

Jaqaman, K., Kuwata, H., Touret, N., Collins, R., Trimble, W.S., Danuser, G., and Grinstein, S. (2011). Cytoskeletal control of CD36 diffusion promotes its receptor and signaling function. Cell 146, 593–606.

Jin, L., Millard, A.C., Wuskell, J.P., Clark, H.A., and Loew, L.M. (2005). Cholesterol-enriched lipid domains can be visualized by di-4-ANEPPDHQ with linear and nonlinear optics. Biophysical journal 89, L04–06.

Johnson, D.I. (1999). Cdc42: An essential Rho-type GTPase controlling eukaryotic cell polarity. Microbiology and molecular biology reviews : MMBR 63, 54–105.

Kalappurakkal, J.M., Anilkumar, A.A., Patra, C., van Zanten, T.S., Sheetz, M.P., and Mayor, S. (2019). Integrin Mechano-chemical Signaling Generates Plasma Membrane Nanodomains that Promote Cell Spreading. Cell.

Kierszniowska, S., Seiwert, B., and Schulze, W.X. (2009). Definition of Arabidopsis sterol-rich membrane microdomains by differential treatment with methyl-beta-cyclodextrin and quantitative proteomics. Molecular & cellular proteomics : MCP 8, 612–623.

Kleine-Vehn, J., Langowski, L., Wisniewska, J., Dhonukshe, P., Brewer, P.B., and Friml, J. (2008). Cellular and molecular requirements for polar PIN targeting and transcytosis in plants. Molecular plant 1, 1056–1066.

Kusumi, A., Koyama-Honda, I., and Suzuki, K. (2004). Molecular dynamics and interactions for creation of stimulation-induced stabilized rafts from small unstable steady-state rafts. Traffic 5, 213–230.

Leyser, O. (2018). Auxin Signaling. Plant physiology 176, 465–479.

Li, C., Yeh, F.L., Cheung, A.Y., Duan, Q., Kita, D., Liu, M.C., Maman, J., Luu, E.J., Wu, B.W., Gates, L., et al. (2015). Glycosylphosphatidylinositol-anchored proteins as chaperones and co-receptors for FERONIA receptor kinase signaling in Arabidopsis. eLife 4.

Li, R., Liu, P., Wan, Y., Chen, T., Wang, Q., Mettbach, U., Baluska, F., Samaj, J., Fang, X., Lucas, W.J., et al. (2012). A membrane microdomain-associated protein, Arabidopsis Flot1, is involved in a clathrin-independent endocytic pathway and is required for seedling development. The Plant cell 24, 2105–2122.

Lin, D., Cao, L., Zhou, Z., Zhu, L., Ehrhardt, D., Yang, Z., and Fu, Y. (2013). Rho GTPase signaling activates microtubule severing to promote microtubule ordering in Arabidopsis. Current biology : CB 23, 290–297.

Lin, W., Tang, W., Anderson, C., and Yang, Z. (2018). FERONIA’s sensing of cell wall pectin activates ROP GTPase signaling in Arabidopsis. bioRxiv.

Lv, X., Jing, Y., Xiao, J., Zhang, Y., Zhu, Y., Julian, R., and Lin, J. (2017). Membrane microdomains and the cytoskeleton constrain AtHIR1 dynamics and facilitate the formation of an AtHIR1-associated immune complex. The Plant journal : for cell and molecular biology 90, 3–16.

Maekawa, M., and Fairn, G.D. (2015). Complementary probes reveal that phosphatidylserine is required for the proper transbilayer distribution of cholesterol. Journal of cell science 128, 1422–1433.

Mahammad, S., and Parmryd, I. (2015). Cholesterol depletion using methyl-beta-cyclodextrin. Methods in molecular biology 1232, 91–102.

Makushok, T., Alves, P., Huisman, S.M., Kijowski, A.R., and Brunner, D. (2016). Sterol-Rich Membrane Domains Define Fission Yeast Cell Polarity. Cell 165, 1182–1196.

Maxwell, K.N., Zhou, Y., and Hancock, J.F. (2018). Rac1 Nanoscale Organization on the Plasma Membrane Is Driven by Lipid Binding Specificity Encoded in the Membrane Anchor. Molecular and cellular biology 38.

Meca, J., Massoni-Laporte, A., Martinez, D., Sartorel, E., Loquet, A., Habenstein, B., and McCusker, D. (2019). Avidity-driven polarity establishment via multivalent lipid-GTPase module interactions. The EMBO journal 38.

Men, S., Boutte, Y., Ikeda, Y., Li, X., Palme, K., Stierhof, Y.D., Hartmann, M.A., Moritz, T., and Grebe, M. (2008). Sterol-dependent endocytosis mediates post-cytokinetic acquisition of PIN2 auxin efflux carrier polarity. Nature cell biology 10, 237–244.

Monshausen, G.B., Miller, N.D., Murphy, A.S., and Gilroy, S. (2011). Dynamics of auxin-dependent Ca2+ and pH signaling in root growth revealed by integrating high-resolution imaging with automated computer vision-based analysis. The Plant journal : for cell and molecular biology 65, 309–318.

Nagawa, S., Xu, T., Lin, D., Dhonukshe, P., Zhang, X., Friml, J., Scheres, B., Fu, Y., and Yang, Z. (2012). ROP GTPase-dependent actin microfilaments promote PIN1 polarization by localized inhibition of clathrin-dependent endocytosis. PLoS biology 10, e1001299.

Nelson, W.J. (2003). Adaptation of core mechanisms to generate cell polarity. Nature 422, 766–774.

Oda, Y., and Fukuda, H. (2012). Initiation of cell wall pattern by a Rho- and microtubule-driven symmetry breaking. Science 337, 1333–1336.

Osher, S., Sethian, J.A. (1988). Fronts propagating with curvature-dependent speed: algorithms based on Hamilton-Jacobi formulations. Journal of Computational Physics 79, 12–49.

Owen, D.M., Rentero, C., Magenau, A., Abu-Siniyeh, A., and Gaus, K. (2012). Quantitative imaging of membrane lipid order in cells and organisms. Nature protocols 7, 24–35.

Platre, M.P., Bayle, V., Armengot, L., Bareille, J., Marques-Bueno, M.D.M., Creff, A., Maneta-Peyret, L., Fiche, J.B., Nollmann, M., Miege, C., et al. (2019). Developmental control of plant Rho GTPase nano-organization by the lipid phosphatidylserine. Science 364, 57–62.

Raghupathy, R., Anilkumar, A.A., Polley, A., Singh, P.P., Yadav, M., Johnson, C., Suryawanshi, S., Saikam, V., Sawant, S.D., Panda, A., et al. (2015). Transbilayer lipid interactions mediate nanoclustering of lipid-anchored proteins. Cell 161, 581–594.

Remorino, A., De Beco, S., Cayrac, F., Di Federico, F., Cornilleau, G., Gautreau, A., Parrini, M.C., Masson, J.B., Dahan, M., and Coppey, M. (2017). Gradients of Rac1 Nanoclusters Support Spatial Patterns of Rac1 Signaling. Cell reports 21, 1922–1935.

Ren, Y., Li, R., Zheng, Y., and Busch, H. (1998). Cloning and characterization of GEF-H1, a microtubule-associated guanine nucleotide exchange factor for Rac and Rho GTPases. The Journal of biological chemistry 273, 34954–34960.

Salehin, M., Bagchi, R., and Estelle, M. (2015). SCFTIR1/AFB-based auxin perception: mechanism and role in plant growth and development. The Plant cell 27, 9–19.

Sartorel, E., Unlu, C., Jose, M., Massoni-Laporte, A., Meca, J., Sibarita, J.B., and McCusker, D. (2018). Phosphatidylserine and GTPase activation control Cdc42 nanoclustering to counter dissipative diffusion. Molecular biology of the cell 29, 1299–1310.

Scheitz, K., Luthen, H., and Schenck, D. (2013). Rapid auxin-induced root growth inhibition requires the TIR and AFB auxin receptors. Planta 238, 1171–1176.

Schrick, K., Mayer, U., Horrichs, A., Kuhnt, C., Bellini, C., Dangl, J., Schmidt, J., and Jurgens, G. (2000). FACKEL is a sterol C-14 reductase required for organized cell division and expansion in Arabidopsis embryogenesis. Genes & development 14, 1471–1484.

Schuck, S., and Simons, K. (2004). Polarized sorting in epithelial cells: raft clustering and the biogenesis of the apical membrane. Journal of cell science 117, 5955–5964.

Sezgin, E., Davis, S.J., and Eggeling, C. (2015). Membrane nanoclusters-tails of the unexpected. Cell 161, 433–434.

Sezgin, E., Levental, I., Mayor, S., and Eggeling, C. (2017). The mystery of membrane organization: composition, regulation and roles of lipid rafts. Nature reviews Molecular cell biology 18, 361–374.

Siegrist, S.E., and Doe, C.Q. (2007). Microtubule-induced cortical cell polarity. Genes & development 21, 483–496.

Simons, K. (2018). Lipid Rafts: A Personal Account. In Physics of Biological Membranes, P. Bassereau, Sens, P., ed. (Cham: Springer).

Simons, K., and Sampaio, J.L. (2011). Membrane organization and lipid rafts. Cold Spring Harbor perspectives in biology 3, a004697.

Sohn, H.W., Tolar, P., Jin, T., and Pierce, S.K. (2006). Fluorescence resonance energy transfer in living cells reveals dynamic membrane changes in the initiation of B cell signaling. Proceedings of the National Academy of Sciences of the United States of America 103, 8143–8148.

Sorek, N., Poraty, L., Sternberg, H., Buriakovsky, E., Bar, E., Lewinsohn, E., and Yalovsky, S. (2017). Corrected and Republished from: Activation Status-Coupled Transient S-Acylation Determines Membrane Partitioning of a Plant Rho-Related GTPase. Molecular and cellular biology 37.

Sorek, N., Segev, O., Gutman, O., Bar, E., Richter, S., Poraty, L., Hirsch, J.A., Henis, Y.I., Lewinsohn, E., Jurgens, G., et al. (2010). An S-acylation switch of conserved G domain cysteines is required for polarity signaling by ROP GTPases. Current biology : CB 20, 914–920.

Souter, M., Topping, J., Pullen, M., Friml, J., Palme, K., Hackett, R., Grierson, D., and Lindsey, K. (2002). hydra Mutants of Arabidopsis are defective in sterol profiles and auxin and ethylene signaling. The Plant cell 14, 1017–1031.

St Johnston, D. (2018). Establishing and transducing cell polarity: common themes and variations. Current opinion in cell biology 51, 33–41.

Stepanova, A.N., Robertson-Hoyt, J., Yun, J., Benavente, L.M., Xie, D.Y., Dolezal, K., Schlereth, A., Jurgens, G., and Alonso, J.M. (2008). TAA1-mediated auxin biosynthesis is essential for hormone crosstalk and plant development. Cell 133, 177–191.

Stone, M.B., Shelby, S.A., Nunez, M.F., Wisser, K., and Veatch, S.L. (2017). Protein sorting by lipid phase-like domains supports emergent signaling function in B lymphocyte plasma membranes. eLife 6.

Strutt, H., and Strutt, D. (2009). Asymmetric localisation of planar polarity proteins: Mechanisms and consequences. Seminars in cell & developmental biology 20, 957–963.

Tan, X., Calderon-Villalobos, L.I., Sharon, M., Zheng, C., Robinson, C.V., Estelle, M., and Zheng, N. (2007). Mechanism of auxin perception by the TIR1 ubiquitin ligase. Nature 446, 640–645.

Varshney, P., Yadav, V., and Saini, N. (2016). Lipid rafts in immune signalling: current progress and future perspective. Immunology.

Vieira, F.S., Correa, G., Einicker-Lamas, M., and Coutinho-Silva, R. (2010). Host-cell lipid rafts: a safe door for micro-organisms? Biology of the cell / under the auspices of the European Cell Biology Organization 102, 391–407.

Vladar, E.K., Antic, D., and Axelrod, J.D. (2009). Planar cell polarity signaling: the developing cell's compass. Cold Spring Harbor perspectives in biology 1, a002964.

Wang, L., Li, H., Lv, X.Q., Chen, T., Li, R.L., Xue, Y.Q., Jiang, J.J., Jin, B., Baluska, F., Samaj, J., et al. (2015). Spatiotemporal Dynamics of the BRI1 Receptor and its Regulation by Membrane Microdomains in Living Arabidopsis Cells. Molecular plant 8, 1334–1349.

Xu, T., Dai, N., Chen, J., Nagawa, S., Cao, M., Li, H., Zhou, Z., Chen, X., De Rycke, R., Rakusova, H., et al. (2014). Cell surface ABP1-TMK auxin-sensing complex activates ROP GTPase signaling. Science 343, 1025–1028.

Xu, T., Wen, M., Nagawa, S., Fu, Y., Chen, J.G., Wu, M.J., Perrot-Rechenmann, C., Friml, J., Jones, A.M., and Yang, Z. (2010). Cell surface- and rho GTPase-based auxin signaling controls cellular interdigitation in Arabidopsis. Cell 143, 99–110.

Yang, Z., and Lavagi, I. (2012). Spatial control of plasma membrane domains: ROP GTPase-based symmetry breaking. Current opinion in plant biology 15, 601–607.

Zhao, X.Y., Li, R.L., Lu, C.F., Baluska, F., and Wan, Y.L. (2015). Di-4-ANEPPDHQ, a fluorescent probe for the visualisation of membrane microdomains in living Arabidopsis thaliana cells. Plant Physiol Bioch 87, 53–60.

Zhou, Y., and Hancock, J.F. (2015). Ras nanoclusters: Versatile lipid-based signaling platforms. Biochimica et biophysica acta 1853, 841–849.

Zhou, Y., Liang, H., Rodkey, T., Ariotti, N., Parton, R.G., and Hancock, J.F. (2014). Signal integration by lipid-mediated spatial cross talk between Ras nanoclusters. Molecular and cellular biology 34, 862–876.

Zhou, Y., Prakash, P., Liang, H., Cho, K.J., Gorfe, A.A., and Hancock, J.F. (2017). Lipid-Sorting Specificity Encoded in K-Ras Membrane Anchor Regulates Signal Output. Cell 168, 239–251 e216.

Zhou, Y., Wong, C.O., Cho, K.J., van der Hoeven, D., Liang, H., Thakur, D.P., Luo, J., Babic, M., Zinsmaier, K.E., Zhu, M.X., et al. (2015). SIGNAL TRANSDUCTION. Membrane potential modulates plasma membrane phospholipid dynamics and K-Ras signaling. Science 349, 873–876.

