## Supplementary Materials for "Auxin-induced nanoclustering of membrane signaling complexes underlies cell polarity establishment in Arabidopsis"

This PDF file includes:

Materials and Methods

Supplementary Text

Figs. S1 to S13

Table S1-S4

Full Reference List

### 1. Materials and Methods

#### Plant Materials and Growth conditions

*A. thaliana* ecotypes *Columbia-0* (*Col-0*) was used as the wild type in this study. The *tmk1tmk4* double mutant and transgenic plants containing 35S::YFP-*CArop6*, 35S::YFP-*DNrop6*, pTMK1::TMK1-GFP or pFER::FER-GFP were described previously (1). The *hyd1-E508* seeds were kindly given by Dr. Kathrin Schrick (Kansas State University, USA). The *fk-J79* seeds were gifts from Dr. Jyan-Chyun Jang (The Ohio State University, USA). The *cpi1-1* seeds were kindly shared by Dr. Markus Grebe (Swedish University of Agricultural Sciences, Sweden). The *wei8-1tar2-1* seeds were kindly shared by Dr. Jaimie Van Norman (University of California, Riverside, USA). The flotillin1-mVenus x *tmk1tmk4* and mEGFP-ROP6 x *tmk1tmk4* mutants were generated by genetic crosses and confirmed by genotyping. Plants were grown in soil or on half-strength Murashige and Skoog agar petri dishes supplemented with 1% (w/v) sucrose. Controlled environmental conditions were provided in the growth room at 22$^{o}$C with 16-hour light/8-hour dark photoperiod.

#### DNA Constructs and Plant Transformation

All constructs were made using the primers listed in Supplemental Table S4. The pGWB601/ROP6::mEGFP-ROP6 plant expression vector was constructed as follows. The *ROP6* promoter (1081 bp upstream of the translational start of *ROP6*) and genomic sequence were amplified from *Col-0* genomic DNA using the following primer pairs: ROP6proF/ROP6proR and ROP6gF/ROP6gR, respectively. The *mEGFP* coding sequence was amplified from the plasmid kindly shared by Dr. Xuemei Chen (University of California Riverside, USA) with the primers mEGFPF and mEGFPR. Those three PCR fragments, *ROP6* promoter, *mEGFP* and *ROP6* genomic sequence, were fused together by the overlapping PCR. The fused PCR products were recombined into the pDONOR207 vector using BP recombination (Invitrogen), and the resulting entry vector was then transferred into Gateway binary vector pGWB601 via the LR reaction. The pCAMBIA1300/Flot1::Flot1-mVenus vector was generated as follows. The 2247-bp promoter region was amplified from *Col-0* genomic DNA using the primers flotmF/flotmR and subcloned into the pCAMBIA 1300 vector via *HindIII* and *BamHI* sites. The genomic sequence (without stop codon) of flotillin 1 was amplified using the primers flotgF/flotgR and the mVenus coding sequence was amplified using the primers mVenusF and mVenusR. Those two PCR fragments were further fused by the overlapping PCR. The fused PCR products were subcloned into the pCAMBIA 1300 vector using *BamHI* and *KpnI* sites. For the 35S::Flot1-mCherry construct, the genomic sequence (without stop codon) of flotillin 1 was amplified using the primers FCF/FCfusR and the mCherry coding sequence was amplified using the primers FCfusF/FCR. Those two PCR fragments were further fused by the overlapping PCR, and then subcloned into the pCAMBIA 1300 vector containing the 35S promoter using *BamHI* and *KpnI* sites. All constructs were sequence verified. Stable transgenic lines were generated by using the standard *Agrobacterium tumefaciens*-mediated floral dip method (*2*) in *Col-0* background.

**IAA, mβCD and oryzalin treatments**

Stock solutions of indole-3-acetic acid (IAA, Sigma-Aldrich) and oryzalin (Sigma-Aldrich) was prepared in dimethyl sulfoxide (DMSO). The stock solution of methyl-β-cyclodextrin (mβCD, Sigma-Aldrich) was prepared in deionized water. A consistent DMSO concentration (0.1%, v/v) was used in all treatments, including mock and mβCD treatments.

For Total Internal Reflection Fluorescence (TIRF) imaging, 2-3 days-old seedlings were incubated in liquid half-strength Murashige and Skoog (½ MS) medium supplemented with IAA, mβCD or oryzalin for the indicated time, then imaged within a 10-min time frame window after the treatments. For IAA treatment alone, the seedlings were incubated in 100nM IAA for 10 min and the mock seedlings were incubated in the medium containing 0.1% DMSO. For the combination of IAA and either oryzalin or mβCD treatments, seedlings were pretreated with 5μM oryzalin or 10mM mβCD for 30mins, and then concomitantly treated with 100nM IAA for 10 min. The mock seedlings were treated with 0.1% DMSO for the same time as the actual treatment.

For pavement cell visualization, seeds were geminated on solid ½ MS medium containing 100nM IAA, 10mM mβCD, 100nM IAA plus 10mM mβCD, or 0.1% DMSO (mock treatment). After germination, liquid ½ MS containing the corresponding chemicals were added onto the solid medium to soak the geminated seeds for additional 2 days. The liquid medium containing the chemicals was refreshed for every 12 hours.

**Confocal laser scanning microscopy analysis of Arabidopsis leaf pavement cell (PC) shape**

For visualizing cotyledon pavement cells in the wild type and mutants, adaxial cotyledon epidermis of 2-3 days-old seedlings were stained in propidium iodide (2mg/ml) solution for 20 mins and then imaged using a Leica TCS SP5 confocal microscopy with the following settings: excitation 535 nm and emission 600-630 nm. The PC images were consistently taken at the mid-region of cotyledons of the same developmental stage. The number of lobes were counted according to the previous method described by Xu et al (2010) (*3*). The indentation width was measured with ImageJ as previously described (*4, 5*).

**ROP6 activity assay**

The ROP6 activity assay was performed as described previously (*6*). Briefly, protoplasts were isolated from leaves of 3-week-old transgenic plants expressing 35S::GFP-ROP6. Isolated protoplasts were treated with half-strength liquid MS media containing100nM IAA for 10 mins or 10mM mβCD for 30 min. Where mβCD were used along with IAA, a pretreatment of 10mM mβCD along was given for 30 min followed by a 10 mins co-treatment with mβCD and 100nM IAA. After treatments, total proteins were extracted from $10^5$ -$10^6$ treated protoplasts using extraction buffer (25 mM HEPES, pH 7.4, 10 mM $MgCl_2$, 10 mM KCl, 5 mM dithiothreitol, 5 mM $Na_3VO_4$, 5 mM NaF,1 mM phenylmethylsulfonyl fluoride, 1% Triton X-100, and protease inhibitor cocktail). Twenty micrograms of MBP-RIC1-conjugated agarose beads was added into the protein extracts and incubated at 4°C for 3hrs. The beads were washed four times by washing buffer at 4°C. GTP-bound ROP proteins associated with the MBP-RIC1 beads were boiled for western blotting with anti-GFP rabbit polyclonal antibodies (Santa Cruz Biotech, Inc) and horseradish peroxidase-conjugated anti-Rabbit IgG antibodies (GE Healthcare). Prior to the pull-down assay, a fraction of total proteins was analyzed by immunoblot assay to determine total ROP6. The experiments were repeated three times.

**ROP6 S-acylation assays**

Assays of ROP6 S-acylation were performed according to the method described by Hemsley et al. (2008)(*7*). IAA treatments were performed on 3-week-old transgenic plants expressing 35S::GFP-ROP6 by spraying. Leaf tissues were collected 10 mins after spraying and immediately flash frozen in liquid nitrogen. The collected samples were ground in liquid nitrogen to a fine powder and then mixed with 1.5ml lysis buffer (100 mM Tris pH 7.2, 150 mM NaCl, 25 mM EDTA, 2.5% SDS, 25mM N-ethylmaleimide) with 1x COMPLETE, Mini EDTA-free Protease Inhibitor Cocktail (Roche). After infiltration through a 200-µm nylon mesh, samples were centrifuged at 16,000 x g for 5 min. The supernatant was collected and protein concentration was determined using the BCA assay. Two mg of proteins were incubated in the lysis buffer for 2hrs at room temperature with gentle mixing and then precipitated using the methanol/chloroform extraction method (*8*). The pellets were resuspended in 1ml of binding buffer (100 mM Tris pH 7.2, 150 mM NaCl, 25 mM EDTA, 2% SDS, 6 M urea with protease inhibitors) and the solution was further divided into two equal aliquots. One aliquot was combined with an equal volume of 1M hydroxylamine. As a control the other aliquot was treated identically but hydroxylamine was replaced with 1M NaCl. After mixing, aliquots of 50 µl was removed and used as a loading control. The remaining samples were incubated with100 µl of a 50% suspension of prewashed Thiopropyl Sepharose CL-6b beads (GE Healthcare Life) at room temperature for 1hr. The beads were washed three times with 1 ml binding buffer. Proteins were eluted by incubating at 37°C for 30 mins in 35 µl 2× SDS sample buffer containing 6M urea and DTT. Samples were analyzed by SDS/PAGE and Western blotting using anti-GFP rabbit polyclonal antibodies (Santa Cruz Biotech, Inc) and horseradish peroxidase-conjugated anti-Rabbit IgG antibodies (GE Healthcare).

**Ratiometric di-4-ANEPPDHHQ fluorescence microscopy imaging of membrane lipid order**

The di-4-ANEPPDHHQ staining was performed according to the procedure previously described (*9-11*). The 2-3 days-old seedlings were incubated in 5 µM di-4-ANEPPDHHQ dissolved in DMSO for 90 mins, washed three times in water, then imaged with a Leica SP5 confocal laser scanning microscopy (CLSM). The live samples were excited with a 488-nm laser, and fluorescence was detected in two emission windows at 500-580 nm and 620-750 nm. Florescence signals were collected by a x40 oil immersion objective. Before the dye was added to the seedlings, no autofluorescence was detected in these two spectral regions when excited with 488-nm laser light. Plasmolysis with 0.8M mannitol revealed that the peripheral staining is associated with plasma membrane, but not with the cell wall (Fig. S1). After the imaging, we followed the published protocol (*11*) to generate pseudo-colored ratiometric general polarization (GP) images. Briefly, the ImageJ macro for GP analysis was applied with the following setting: the threshold value was fixed at 15, the color scale for the output GP images was set to 'grays', and no immunofluorescence mask was selected. The GP values were calculated according to the following equations:

$$\mathrm{GP} = (I_{500\text{-}580} - GI_{620\text{-}750}) / (I_{500\text{-}580} + GI_{620\text{-}750}) \qquad (1)$$

$$G = (\mathrm{GP}_{ref} + \mathrm{GP}_{ref}\,\mathrm{GP}_{mes} - \mathrm{GP}_{mes} - 1) / (\mathrm{GP}_{mes} + \mathrm{GP}_{ref}\,\mathrm{GP}_{mes} - \mathrm{GP}_{ref} - 1) \qquad (2)$$

Where *I* represents the intensity in each pixel in the images collected from two spectral channels of CLSM, 500–580 nm and 620–750 nm; G is a calibration factor. It is used to compensate for different collection efficiency between two channels and is calculated according to equation (2); $\mathrm{GP}_{mes}$ is the GP value of di-4-ANEPPDHQ in pure DMSO solution with the same microscopy settings as those used for the real sample. It is calculated using equation (1) with G=1; $\mathrm{GP}_{ref}$ is a reference value for di-4-ANEPPDHQ in DMSO, here fixed at -0.85 as suggested (*11*).

Based on the generated GP images, we further extracted the mean intensity values at the complementary lobing and indenting sides of the plasma membranes by employing the polygon selection and measurement tools in ImageJ. The extracted values were further normalized with the following equation: GP = (extracted values/127.5) – 1.

**TIRF imaging**

Cotyledons of 2-3 days-old seedlings were imaged on a Zeiss Elyra PS1 system (Zeiss, Germany) with a 100 × Apo (numerical aperture 1.46) oil-immersion objective, in Total Internal Reflection Fluorescence (TIRF) mode. The optimum critical angle was determined as giving the best signal-to-noise ratio. Pixel size was 0.107 µm. The development of Arabidopsis leaf pavement cells is separated into three stages (*4, 12*). To minimize developmental differences, stage II cells with shallow lobes were selected for imaging and the focus was set to the mid-region of pavement cells. Movies were recorded at a rate of 10 frames per second (100ms exposure time) on a 256 x 256-pixels region of interest for 200 frames. GFP was excited using a 488 nm solid-state laser diode and the emission was collected with an EM-CCD camera with bandwidth filters ranging from 495–550 nm. For two-color TIRF, the ONI Nanoimager (Oxford Nanoimaging Ltd, Oxford, UK) was used. GFP- and mCherry-labeled proteins were exited using 488 nm and 561 nm laser lines, respectively. Fluorescence emission was collected with bandwidth filters ranging from 495–550 nm for GFP and 570–640 nm for mCherry.

#### Spot size measurement and single-particle tracking

Particle tracking and trajectory analysis was performed using a custom MATLAB R2018a extension code with the Spots Detection module in Imaris (Bitplane) software. Prior to spot estimation background subtraction was applied. Estimated spots were modeled with a minimal XY diameter spot size set to 0.3 µm, followed by further filtering using the default "quality" setting automatically calculated by Imaris. Filtered spots were served as seed points for calculating spot sizes using a region growing method. The spot region growth was halted when the local contrast around the border of the spot region met the automatic threshold calculated by Imaris. Spot diameters were then recalculated based on the region volume detected in the previous step. No manual editing was performed.

An autoregressive motion algorithm was used for particle tracking. A maximum distance was set to 0.5 µm. This setting disallows connections between a spot and a candidate match if the distance between the predicted future position of the spot and the candidate position exceeds 0.5 µm. A gap-closing algorithm was used to compensate undetected spots in one or more of the consecutive frames along a spot trajectory to prevent inaccurately fragmented tracks. A maximum gap size, i.e., the allowed maximum number of consecutive time frames with missing spots, was set to 8. Single-particle tracking trajectories in Supplementary Videos 1, 2, 4 and 5 are shown as dragontail visualization with 10 time points of the track length.

#### Trajectory Analysis

The mean-square displacement (MSD) and diffusion coefficient were analyzed following a method developed earlier (*13*). Trajectories with length > 8 frames were selected for MSD and diffusion coefficient analysis. For each track, the MSD (*t*) was calculated from the following formula:

$$\mathrm{MSD}(t) = \frac{1}{L-n}\sum_{s=0}^{l-n-1}\left(r(s+n) - r(s)\right)^2,$$

Where $n = t/\Delta t$, $L$ is the length of the trajectory (number of frames), and r(*s*) is the two-dimensional position of the particle in frame *s* ($s = 0$ corresponds to the start of the trajectory). The diffusion coefficient for a spot was determined by fitting a line to MSD ($n\Delta t$) with *n* ranging from 1 to the largest integer $\leq L/4$ (*13*).

The natural logarithm of diffusion coefficients for all tracks (>8 frames) in each condition were pooled to obtain the log(D) density histogram with the number of bins of 800. The data was then fitted into one-component and two-component Gaussian mixture (univariate, unequal variance) models by using the R mClust Package with the default settings. The package provides the Bayesian Information Criterion (BIC) to assess how well the model explains the data. For all density distributions of TMK1, Flotillin1 and ROP6 that were obtained and analyzed in this study, the two-component model yielded significantly higher BIC values than the one-component model, indicating the better fit. Each fit to the two-component model outputs a set of parameters: the mean ($\mu_1$, $\mu_2$), standard deviation ($\sigma_1$, $\sigma_1$) and proportional weights ($\alpha_1$, $\alpha_2$) for each component in the model. The mean ($m_1$,$m_2$) and variance ($v_1$, $v_2$) of the lognormal distribution for each component is calculated based on the following equations:

$m$=exp($\mu$+$\sigma^2$/2)

$v$=exp(2$\mu$+$\sigma^2$)(exp($\sigma^2$)−1)

The overall average diffusion coefficient ($m_{overall}$) and variance ($v_{overall}$) is calculated based on the following equations:

$m_{overall} = m_1 * \alpha_1 + m_2 * \alpha_2$

$v_{overall} = \alpha_1 * v_1 + \alpha_1 *( m_1 - m_{overall})^2 + \alpha_2 * v_2 + \alpha_2 *( m_2 - m_{overall})^2$

**Statistical Analyses**

All data analyses were performed with PRISM software (GraphPad Software Inc., La Jolla, CA). Each dataset was subjected to the D'Agostino & Pearson normality test to assess normality of the residuals. Based on the p-value obtained from the normality test, parametric ($p \geq 0.05$) or non-parametric tests ($p < 0.05$) were performed. For parametric test, statistical analyses between two groups were assessed by using the unpaired t test with Welch's correction. For non-parametric test, the Mann-Whitney U test was used to compare differences between two groups. ($*P < 0.05$, $** \leq 0.01$, $***P \leq 0.001$ and $****$ $P \leq 0.0001$).

**Mathematical modeling and simulation**

We model the process of early polarity establishment in pavement cells by a set of partial differential equations on a closed moving curve in two dimensions. The system of equations considers the concentration of auxin (aux), TMK1 nanoclusters, ordered lipid nanodomains, ROP6 nanoclusters and cortical microtubules (CMTs). TMK1 nanoclusters, ordered lipid nanodomains and ROP6 nanoclusters are modeled as diffusive molecules on the plasma membrane. Auxin effect is modeled by proportionally increasing the initial concentration of TMK1 nanoclusters and ordered lipid nanodomains, as our experimental data show that auxin promotes TMK1 nanoclustering and lipid ordering. The governing equations of the dynamics are shown as below (for simplicity, we use Lipid instead of ordered lipid nanodomain and MT for CMT in the following equations):

$$\frac{\partial[TMK]}{\partial t}=\nabla_s\cdot\left(D_T\nabla_s[TMK]\right)-d_T[TMK]$$

$$\frac{\partial[Lipid]}{\partial t}=\nabla_s\cdot\left(D_L\nabla_s[Lipid]\right)-d_L[Lipid]$$

$$\frac{\partial[ROP]}{\partial t}=\nabla_s\cdot\left(D_R\nabla_s[ROP]\right)-d_R[ROP]+R_{\min}+\frac{R_{\max}-R_{\min}}{1+\left(\frac{[TMK][Lipid]}{k_{TLR}}\right)^{-n_{TLR}}}$$

$$\frac{\partial[MT]}{\partial t}=T_{\min}+\frac{T_{\max}-T_{\min}}{1+\left(\frac{[ROP]}{k_{RM}}\right)^{-n_{RM}}}-d_M[MT]$$

In particular, the associated diffusion is nonhomogeneous and the rates are regulated as below:

$$D_T=\frac{\overline{D_T}}{1+\left(\frac{[Lipid]}{k_{LT}}\right)^{n_{LT}}+\left(\frac{[MT]}{k_{MT}}\right)^{n_{MT}}}$$

$$D_L=\frac{\overline{D_L}}{1+\left(\frac{[TMK]}{k_{TL}}\right)^{n_{TL}}+\left(\frac{[MT]}{k_{ML}}\right)^{n_{ML}}}$$

$$D_R=\frac{\overline{D_R}}{1+\left(\frac{[Lipid]}{k_{LR}}\right)^{n_{LR}}}$$

Here, the notation $\nabla_S$ denotes the gradient over the interface, and $\nabla_S\cdot(D_*\nabla_S)$ represents the surface diffusion along the cell membrane with $D_*$ denoting the diffusion coefficient. The parameter $d_*$ denotes the degradation rate. The coefficients $R_{min}$ and $R_{max}$ represent the minimal and maximal synthesis rate of ROP6, respectively, so do the coefficients $T_{min}$ and $T_{max}$ for CMTs. Since TMK1 nanoclusters and order lipid nanodomains restrict the diffusion of each other, the diffusion coefficients are negatively regulated by each other in the model. ROP6 is activated at sites where both TMK1 nanoclusters and ordered lipid nanodomains have high concentration, which is modeled by a Hill function in terms of the product of concentrations of TMK1 nanoclusters and ordered lipid nanodomains. Moreover, ROP6 diffusion is restricted by ordered lipid nanodomains. ROP6 signaling controls CMT ordering, which further restricts the diffusion of TMK1 nanoclusters and ordered lipid nanodomains as a feedback. Both positive and negative regulations are modeled by Hill functions with $k_*$ denoting the EC50 values and $n_*$ for Hill coefficients.

The closed moving interface representing cell membrane is computed by using level set method (*14*). In particular, a level set function $\phi$ is defined on a two-dimensional domain in rectangular shape which is sufficiently large to cover the entire cell throughout the time of interest. The level set function $\phi$ satisfies that it is always negative inside the cell and positive outside the cell. In this study, we use the signed distance function for $\phi$. The membrane can be captured by extracting the zero contour of $\phi$. Therefore, instead of computing the moving interface explicitly, we solve a Hamilton-Jacobian equation for $\phi$ which describes the morphological change of cell membrane and obtains the moving interface implicitly on a fixed Cartesian mesh. The governing equation for $\phi$ is

$$\phi_t + \boldsymbol{u}\cdot\nabla\phi = 0, \tag{1}$$

where u denotes the corresponding velocity field of the cell shape change. In this study, we assume the cell undergoes an isotropic growth due to some other signals which are not modeled specifically in this network. In addition to that, CMTs provide a resistance force to prevent the cell from expansion. Therefore, the velocity field is assumed to be

$$\boldsymbol{u} = \max\big(0, a - k[MT]([MT]\geq\theta)\big)\vec{n}. \tag{2}$$

Here, $a$ is the magnitude of the isotropic growth. The resistance force exerted by CMTs leads to a negative velocity field that is linearly dependent on the concentration of CMTs with coefficient $k$. The parameter $\theta$ is the minimal value such that the resistance force exerted by CMTs becomes effective. The vector $\vec{n}$ denotes the normal direction to the moving interface.

After taking into account the deformation of cell membrane, the dynamics of each surfactant associated with the membrane can be described by a convection-diffusion-reaction equation on the moving interface in the form of

$$\frac{\partial[TMK]}{\partial t} + u\cdot\nabla[TMK] - \vec{n}\cdot\nabla u\cdot\vec{n}[TMK] = \nabla_s\cdot\big(D_T\nabla_s[TMK]\big) - d_T[TMK]$$

$$\frac{\partial[Lipid]}{\partial t} + u\cdot\nabla[Lipid] - \vec{n}\cdot\nabla u\cdot\vec{n}[Lipid] = \nabla_s\cdot\big(D_L\nabla_s[Lipid]\big) - d_L[Lipid]$$

$$\frac{\partial[ROP]}{\partial t} + u\cdot\nabla[ROP] - \vec{n}\cdot\nabla u\cdot\vec{n}[ROP] = \nabla_s\cdot\big(D_R\nabla_s[ROP]\big) - d_R[ROP] + R_{\min} + \frac{R_{\max} - R_{\min}}{1 + \left(\frac{[TMK][Lipid]}{k_{TLR}}\right)^{-n_{TLR}}}$$

$$\frac{\partial[MT]}{\partial t} + u\cdot\nabla[MT] - \vec{n}\cdot\nabla u\cdot\vec{n}[MT] = T_{\min} + \frac{T_{\max} - T_{\min}}{1 + \left(\frac{[ROP]}{k_{RM}}\right)^{-n_{RM}}} - d_M[MT]$$

where $\vec{n}$ is the normal vector of the moving interface. The extra two terms on the left-hand side represents the advection and the dilution of the surfactant on the interface due to the morphological change. Therefore, the entire system includes equations of TMK nanoclusters, ordered lipid nanodomains, ROP and CMTs, together with the equation of $\phi$ to be solved simultaneously.

To compute the surface diffusion of a surfactant $f$ numerically, the nonhomogeneous surface Laplacian is decomposed as below:

$$\nabla_s \cdot \left(D(x)\nabla_s f\right) = \nabla_s D(x)\cdot \nabla_s f + D(x)\nabla_s \cdot \nabla_s f$$
$$= \nabla_s D(x)\cdot \nabla_s f + D(x)\left(\Delta f - \frac{\partial^2 f}{\partial n^2} - \kappa\frac{\partial f}{\partial n}\right)$$
$$= \nabla D(x)\cdot \nabla f - \frac{\partial D(x)}{\partial n}\frac{\partial f}{\partial n} + D(x)\left(\Delta f - \frac{\partial^2 f}{\partial n^2} - \kappa\frac{\partial f}{\partial n}\right)$$

where $\Delta$ is the regular two-dimensional diffusion operator and $\kappa$ is the curvature of the moving interface which can be calculated in terms of $\phi$ as $\kappa = \nabla \cdot \left(\frac{\nabla\phi}{|\nabla\phi|}\right)$. Notice that this decomposition requires the variable $f$ defined in the 2d domain instead of the moving interface only. Therefore, we perform an extension of $f$ such that it remains constant along the normal direction, by solving the following equation until its steady state.

$$f_t + \mathrm{sgn}(\phi)\vec{n}\cdot\nabla f = 0, \tag{3}$$

where the function *sgn(x)* denotes the sign of *x*, i.e., *sgn(x)=-1* if *x<0* and *sgn(x)=1* if *x>0*. Therefore the equations for the extended variables are shown as below

$$\frac{\partial[TMK]}{\partial t} + \left(u - \nabla D_T\right)\cdot\nabla[TMK] - \vec{n}\cdot\nabla u\cdot\vec{n}[TMK] = D_T\left(\Delta[TMK] - \frac{\partial^2[TMK]}{\partial n^2} - \kappa\frac{\partial[TMK]}{\partial n}\right) - \frac{\partial D_T}{\partial n}\frac{\partial[TMK]}{\partial n} - d_T[TMK]$$
$$\frac{\partial[Lipid]}{\partial t} + \left(u - \nabla D_L\right)\cdot\nabla[Lipid] - \vec{n}\cdot\nabla u\cdot\vec{n}[Lipid] = D_L\left(\Delta[Lipid] - \frac{\partial^2[Lipid]}{\partial n^2} - \kappa\frac{\partial[Lipid]}{\partial n}\right) - \frac{\partial D_L}{\partial n}\frac{\partial[Lipid]}{\partial n} - d_L[Lipid]$$
$$\frac{\partial[ROP]}{\partial t} + \left(u - \nabla D_R\right)\cdot\nabla[ROP] - \vec{n}\cdot\nabla u\cdot\vec{n}[ROP] = D_R\left(\Delta[ROP] - \frac{\partial^2[ROP]}{\partial n^2} - \kappa\frac{\partial[ROP]}{\partial n}\right) - \frac{\partial D_R}{\partial n}\frac{\partial[ROP]}{\partial n} - d_R[ROP] + R_{\min} + \frac{R_{\max} - R_{\min}}{1 + \left(\frac{[TMK][Lipid]}{k_{TLR}}\right)^{-n_{TLR}}}$$
$$\frac{\partial[MT]}{\partial t} + u\cdot\nabla[MT] - \vec{n}\cdot\nabla u\cdot\vec{n}[MT] = T_{\min} + \frac{T_{\max} - T_{\min}}{1 + \left(\frac{[ROP]}{k_{RM}}\right)^{-n_{RM}}} - d_M[MT] \tag{4}$$

The numerical algorithm is summarized in the following:

**Step 0**: Set up the initial condition for $\phi$ representing initial cell shape, TMK nanoclusters, ordered lipid nanodomains, ROP nanoclusters and CMTs.

**Step 1**: Extend the variables TMK nanoclusters, ordered lipid nanodomains, ROP nanoclusters and CMTs along with the current $\phi$ according to (3).

**Step 2**: Calculate the velocity field according to (2)

**Step 3**: Evolve (1) and (4) by using the velocity field obtained in Step 2 by one time step.

**Step 4**: Reinitialize the level set function $\phi$ to be the signed distance function if necessary. Go back to repeat Step 1.

In **Step 0**, for the initial condition, the cell shape is modeled by using a rounded rectangle with aspect ratio 4:5 to match the experimental observation. The initial concentration of TMK nanoclusters and ordered lipid nanodomains are provided along the cell membrane uniformly with some random multiplicative noise added as shown below

$$TMK = \mu_{TMK} + \alpha\sqrt{\mu_{TMK}}X_1(x,y), \qquad Lipid = \mu_{Lipid} + \alpha\sqrt{\mu_{Lipid}}X_2(x,y),$$

where $\mu_{\cdot}$ represents the mean concentration and $\alpha$ denotes the noise amplitude. In all simulations we choose $\alpha = 0.3$ to include significant noise into the dynamics. $X_{\cdot}(x,y)$ is a random variable following standard normal distribution. Notice that auxin treatment leads to a proportional increase in the initial concentration of TMK1 nanoclusters and ordered lipid nanodomains (the ratio of the number of TMK1 nanoclusters with size > 300 nm in the auxin treatment to that in the mock treatment is 1.068 in our experiment), while mβCD treatment leads to a reduction in the initial concentration of ordered lipid nanodomains only (the ratio of the number of ordered lipid nanodomains with size > 350 nm in the mock treatment to that in the mβCD treatment is 1.0532 in our experiments). Therefore, we set the mean of the TMK1 concentration to be 1 and ordered lipid nanodomains to be 1.4 in the control condition, and set the mean concentration under other conditions proportionally. The details are provided in the following table.

| Mean concentration | TMK1 nanoclusters | Ordered lipid nanodomains |
|---|---|---|
| Ctr | 1 | 1.4 |
| mβCD | 1 | 1.2 |
| Auxin | 1.1 | 1.4×1.1 |
| Auxin+mβCD | 1.1 | 1.2×1.1 |

In simulations, the fold change in the initial mean concentration of TMK nanoclusters and ordered lipid nanodomains between Auxin/Ctr=1.1; the fold change in the initial mean concentration of ordered lipid nanodomains between Ctr/mβCD=1.167, which are close to the experimental data. The initial profiles of the concentration of TMK nanoclusters and ordered lipid nanodomains for one typical sample in wild type are shown as below.

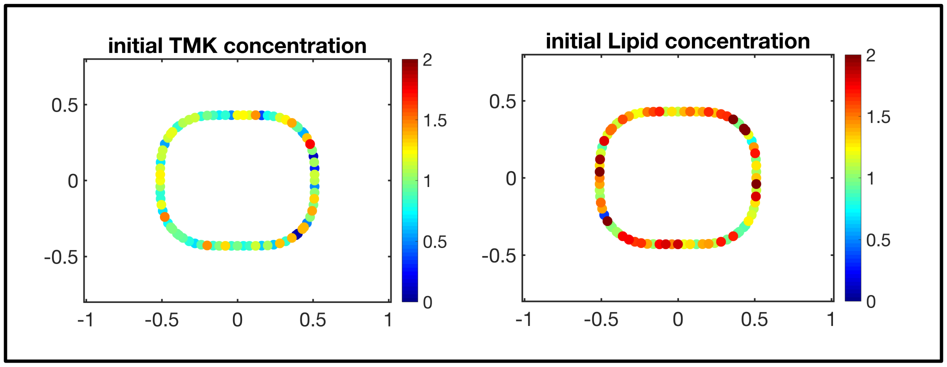

The numerical algorithm we used to compute the model was developed in (*15*). The computation was performed in MATLAB with spatial mesh size $h = 0.04\mu m$ and temporal step size $dt = 0.1s$

to guarantee the stability and accuracy. Each simulation runs until cell area reaches a 1.5-fold increase.

**Parameters used in the mathematical models unless otherwise specified**

| Parameter | Simulation | Experiment | Parameter | Simulation | Experiment |
|---|---|---|---|---|---|
| **Diffusion Coefficient** | $\mu m^2/s$ | $\mu m^2/s$ | **Degradation rate** | $s^{-1}$ | |
| $\boldsymbol{D_T}$ | 0.01 | 0.150±0.001 | $d_T$ | 0.0001 | - |
| $\boldsymbol{D_L}$ | 0.01 | 0.052±0.004 | $d_L$ | 0.0001 | - |
| $\boldsymbol{D_R}$ | 0.01 | 0.162±0.003 | $d_R$ | 0.5 | - |
| | | | $d_M$ | 0.01 | - |
| **Production Rate** | $s^{-1}$ | | **Cell Deformation** | $s^{-1}$ | |
| ***Rop_min*** | 0 | - | $\alpha$ | $5\times10^{-5}$ | - |
| ***Rop_max*** | 1.3 | - | $k$ | $5\times10^{-5}$ | - |
| ***MT_min*** | 0 | - | $\theta$ | 0.01 | - |
| ***MT_max*** | 0.04 | - | | | |
| **EC50** | | | **Hill Coefficient** | | |
| **kLT** | 0.2 | - | nLT | 2 | - |
| **kTL** | 0.2 | - | nTL | 2 | - |
| **kLR** | 0.005 | - | nLR | 10 | - |
| **kRM** | 1.3 | - | nRM | 4 | - |
| **kMT** | 0.006 | - | nMT | 6 | - |
| **kML** | 0.006 | - | nML | 6 | - |
| **kTLR** | 1.3 | - | nTLR | 4 | - |

**Data availability**

All data that support the findings of this study are available from the corresponding author upon request.

Table S1. Summary of diffusion characteristics of TMK1 particles. Data were presented as mean ± SEM.

| | Treatment | # of tracks | Population 1 (%) | Population 2 (%) | $D_0$ ($\mu m^2/s$) | $D_1$ ($\mu m^2/s$) | Average D ($\mu m^2/s$) |
|---|---|---|---|---|---|---|---|
| TMK1 in Col | DMSO | 156480 | 51.34% | 48.66% | 0.150±0.001 | 0.391±0.002 | 0.267±0.001 |
| TMK1 in Col | IAA | 131187 | 52.16% | 47.84% | 0.132±0.001 | 0.369±0.002 | 0.245±0.001 |
| TMK1 in Col | Oryzalin | 85436 | 47.74% | 52.26% | 0.227±0.003 | 0.485±0.003 | 0.362±0.002 |
| TMK1 in Col | Oryzalin + IAA | 96457 | 50.93% | 49.07% | 0.153±0.002 | 0.394±0.002 | 0.271±0.001 |
| TMK1 in Col | mβCD | 98267 | 48.76% | 51.24% | 0.249±0.003 | 0.537±0.003 | 0.396±0.002 |
| TMK1 in Col | mβCD + IAA | 107269 | 49.08% | 50.92% | 0.244±0.003 | 0.545±0.003 | 0.397±0.002 |

Table S2. Summary of diffusion characteristics of flotillin1(Flot) particles. Data were presented as mean ± SEM

| | Treatment | # of tracks | Population 1 (%) | Population 2 (%) | $D_0$ ($\mu m^2/s$) | $D_1$ ($\mu m^2/s$) | Average D ($\mu m^2/s$) |
|---|---|---|---|---|---|---|---|
| Data presented in Figure 4 | | | | | | | |
| Flot in Col | DMSO | 49216 | 56.42 | 43.58 | 0.052 ±0.004 | 0.192±0.002 | 0.113±0.002 |
| Flot in Col | IAA | 43223 | 65.74 | 34.26 | 0.036±0.003 | 0.129±0.002 | 0.068±0.002 |
| Flot in *tmk1tmk4* | DMSO | 47078 | 34.73 | 65.27 | 0.374±0.039 | 0.831±0.006 | 0.672±0.012 |
| Flot in *tmk1tmk4* | IAA | 48206 | 33.93 | 66.07 | 0.396±0.052 | 0.848±0.007 | 0.695±0.014 |
| Data presented in Figure 7 | | | | | | | |
| Flot in Col | DMSO | 36937 | 51.28 | 48.72 | 0.065±0.006 | 0.154±0.003 | 0.109±0.003 |
| Flot in Col | IAA | 47316 | 52.52 | 47.48 | 0.025±0.002 | 0.118±0.002 | 0.069±0.002 |
| Flot in Col | Oryzalin | 53899 | 50.95 | 49.05 | 0.078±0.007 | 0.182±0.002 | 0.129±0.003 |
| Flot in Col | Oryzalin + IAA | 50214 | 52.59 | 47.41 | 0.055±0.005 | 0.140±0.002 | 0.095±0.003 |

Table S3. Summary of diffusion characteristics for ROP6 particles. Data were presented as mean ± SEM.

| | Treatment | # of tracks | Population 1 (%) | Population 2 (%) | $D_0$ ($\mu m^2/s$) | $D_1$ ($\mu m^2/s$) | Average D |
|---|---|---|---|---|---|---|---|
| *CArop6* | DMSO | 86712 | 35.30 | 64.70 | 0.162±0.003 | 0.260±0.002 | 0.225±0.002 |
| *DNrop6* | DMSO | 97390 | 28.93 | 71.07 | 0.166±0.003 | 0.299±0.001 | 0.261±0.001 |
| ROP6 (wt) in Col | DMSO | 40685 | 16.20 | 83.80 | 0.259±0.009 | 0.422±0.002 | 0.396±0.002 |
| ROP6 (wt) in Col | Auxin | 42016 | 17.70 | 82.30 | 0.248±0.008 | 0.395±0.002 | 0.369±0.002 |
| ROP6 (wt) in *tmk1tmk4* | DMSO | 32663 | 15.32 | 84.68 | 0.372±0.013 | 0.636±0.004 | 0.596±0.004 |
| ROP6 (wt) in *tmk1tmk4* | Auxin | 32893 | 15.68 | 84.32 | 0.378±0.011 | 0.652±0.004 | 0.609±0.004 |

Table S4. Primers used for vector construction

| Primers name | Primer sequence |
|---|---|
| ROP6-ProF | GGGGACAAGTTTGTACAAAAAAGCAGGCTCCCAAGCTTTCAGAAAAGAGGATGATATAAG |
| ROP6-ProR | CTCGCCCTTGCTCACCATTCTAGACTTTCTCTCCTTCTTCAAACTTCAAAAACC |
| ROP6-gF | CATGGACGAGCTGTACAAGGGCGGAGGAGGATCCATGAGTGCTTCAAGGTTTATCAAGTG |
| ROP6-gR | GGGGACCACTTTGTACAAGAAAGCTGGGTCTCAGAGTATAGAACAACCTTTCTGAGATTT |
| mEGFP-F | GGTTTTTGAAGTTTGAAGAAGGAGAGAAAGTCTAGAATGGTGAGCAAGGGCGAG |
| mEGFP-R | CACTTGATAAACCTTGAAGCACTCATGGATCCTCCTCCGCCCTTGTACAGCTCGTCCATG |
| FlotmF | TATAAAGCTTGAACTGGTTGTTGATGATGATGATG |
| FlotmR | TATAGGATCCCTTTTGATTTAATTTGGACTTTTCGCC |
| flotgF | TATAGGATCCATGTTCAAAGTTGCAAGAGCGTCAC |
| flotgR | GCTCCTCGCCCTTGCTCACCATAGATCCTCCTCCGCCGCTGCGAGTCACTTGCTTCGG |
| mVenusF | CCGAAGCAAGTGACTCGCAGCGGCGGAGGAGGATCTATGGTGAGCAAGGGCGAGGAGC |
| mVenusR | TATAGGTACCTTACTTGTACAGCTCGTCCATGCCG |
| FCF | TATAGGATCCATGTTCAAAGTTGCAAGAGCGTCAC |
| FCfusR | TCCTCCTCGCCCTTGCTCACCATTCTAGAGGATCCGCTGCGAGTCACTTGCTTCGGTTCC |
| FCfusF | CCGAAGCAAGTGACTCGCAGCGGATCCTCTAGAATGGTGAGCAAGGGCGAGGAGGATAAC |
| FCR | TATAGGTACCTTACTTGTACAGCTCGTCCATGCCGCC |

2. **Author Contribution Statement**

The project was conceived and supervised by Z.Y. W.C. developed the mathematical models. X.P. conceived the study, designed and performed all experiments except for ROP activity assays. L.F. and U.M. provided the MATLAB code for single-particle tracking. J.L. developed the Python code for batch data processing. J.L. and B.S.A. developed the MATLAB code for diffusion coefficient analyses. W.L. conducted ROP activation assays. S.Z. assisted in the genotyping experiments. T.Z. assisted in the TIRF imaging. Z.Y., W.C. and X.P. wrote the manuscript.

Supplementary Video 1: Example trajectories of Flot1-mVenus on a 2-3 days-old Arabidopsis cotyledon pavement cell. The movie was acquired at 10 frames per second (100ms exposure time) on a 256 x 256-pixels region of interest for 200 frames using TIRF microscopy. The dynamic behavior of Flot1-mVenus particles was tracked by dragontrail command of IMARIS Track, in which 10 time points of the track length were shown. Trajectories are shown with color coding for absolute time to visualize the progressive dynamics. Scale bar, 2µm.

Supplementary Video 2: Dynamic assembly and disassembly of Flot1-mVenus particles on Arabidopsis cotyledon pavement cells. The movie was acquired at 10 frames per second (100ms exposure time) for 100 frames using TIRF microscopy. **Left Video**: Raw video used for size measurement and single-particle tracking. **Middle Video:** Dynamics size-measurement of Flot1-mVenus particles. The size of Flot1-mVenus particles was measured using a region growing method in IMARIS. **Right Video:** Trajectories of Flot1-mVenus particles. The dynamic behavior of Flot1-mVenus particles was tracked by dragontrail command of IMARIS Track, in which 10 time points of the track length were shown. Trajectories are shown with color coding for absolute time to visualize the progressive dynamics. Scale bar, 0.2µm.

Supplementary Video 3: 3D representation of a pavement cell expressing Flot1-mCherry driven by the (CaMV) 35S promoter. After *Agrobacterium*-mediated floral-dip transformation, we obtained a stable transgenic line with chimeric expression of the flot-mCherry transgene. The sporadic expression of flot-mCherry allows us to visualize the distribution of flotilllin1 proteins at the plasma membrane of single cells. The 3D image was reconstructed from 43 optical sections spaced at 0.55µm increments. Scale bar, 20 µm.

Supplementary Video 4: Example trajectories of TMK1-GFP on a 2-3 days-old Arabidopsis cotyledon pavement cell. The movie was acquired at 10 frames per second (100ms exposure time) on a 256 x 256-pixels region of interest for 200 frames using TIRF microscopy. The dynamic behavior of TMK1-GFP particles was tracked by dragontrail command of IMARIS Track, in which 10 time points of the track length were shown. Trajectories are shown with color coding for absolute time to visualize the progressive dynamics. Scale bar, 2µm.

Supplementary Video 5: Example trajectories of mEGFP-ROP6 on a 2-3 days-old Arabidopsis cotyledon pavement cell. The movie was acquired at 10 frames per second (100ms exposure time) on a 256 x 256-pixels region of interest for 200 frames using TIRF microscopy. The dynamic behavior of mEGFP-ROP6 particles was tracked by dragontrail command of IMARIS

Track, in which 10 time points of the track length were shown. Trajectories are shown with color coding for absolute time to visualize the progressive dynamics. Scale bar, 2μm.

Supplementary Video 6: Representative simulation showing the process of multi-polarity formation in the pavement cell. The color scale indicates the level of active ROP6.

Supplementary Video 7: Representative simulation showing that when the auxin concentration is higher than the optimum level, a low degree of lateral segregation of active ROP6 domains results in less polarity sites. The color scale indicates the level of active ROP6.

3. **Supplemental Figures and Legends**

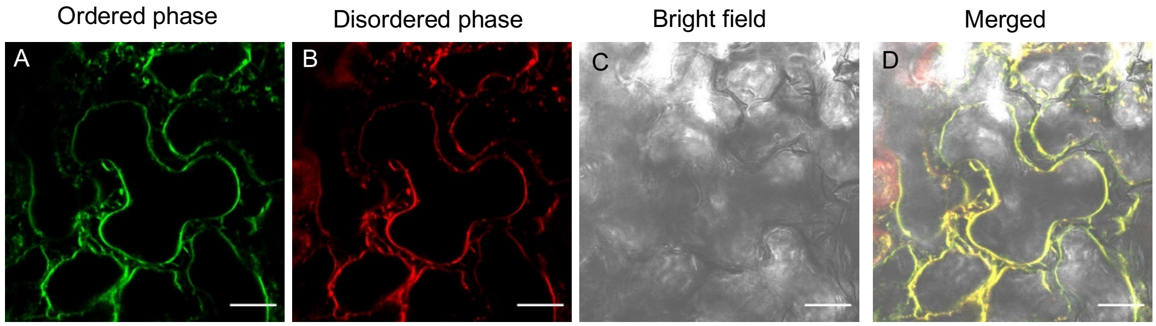

**fig S1. Di-4-ANEPPDHQ does not label the cell wall of epidermal pavement cells.** Representative images obtained from 90 mins incubation of 2-3 days-old cotyledons with di-4-ANEPPDHHQ followed by plasmolysis with 0.8 M mannitol for 5mins. (A) shows di-4-ANEPPDHHQ fluorescence recorded between 500-580 nm. (B) shows ANEPPDHHQ fluorescence recorded between 620-750nm. (C) shows the bright-field image. (D) shows the merged image.  Scale bars = 15μm.

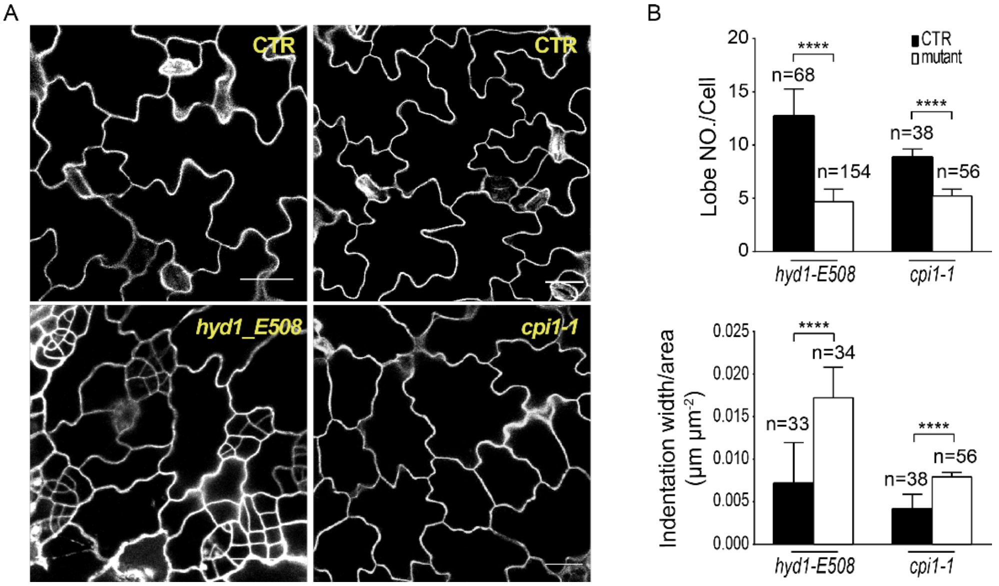

**fig S2. Sterol biosynthesis mutants show altered pavement cell shape. (A)** Representative confocal images of pavement cells in cotyledons of sterol biosynthesis mutants and their corresponding wild-types. *HYDRA1* gene encodes a sterol Δ8-Δ7 sterol isomerase (*16, 17*) and FRACKEL gene encodes a sterol C14 reductase (*18*). The seedlings of *hyd1* and *fk-J79* mutants are similarly defective in bulk sterol biosynthesis, with reduced levels of campesterol and sitosterol but increased stigmasterol compared with wild type (*16*). Cyclopropylsterol isomerase1 (CPI1) functions to open the cyclopropyl ring of cycloeucalenol to produce obtusifoliol, the step in which the *cpi1-1* mutant is defective (*19*). Homozygous *cpi1-1* plants accumulate cyclopropylsterols up to 99% of its total sterol content with an almost complete reduction of wild-type sterols, including sitosterol, stigmasterol, campesterol. All these homozygous mutants display near-sterile or sterile phenotype. The heterozygous mutants were segregated into wild-type (top panel) and homozygous mutants (bottom panel). The pavement cell phenotype was analyzed at 7days after germination. **(B, top panel)** Quantitative analysis of the number of lobes for wild-type and mutants. The average lobe number per cell in mutants was fewer than the wild type. **(B, bottom panel)** Quantitative analysis of indentation widths for wild-type and mutants. The average indentation width of mutants was significantly increased compared with that of the wild type. *n* represents the number of independent cells. Scale bars = 30 μm. Data were presented as mean ± SD. See methods for detailed statistical analyses.

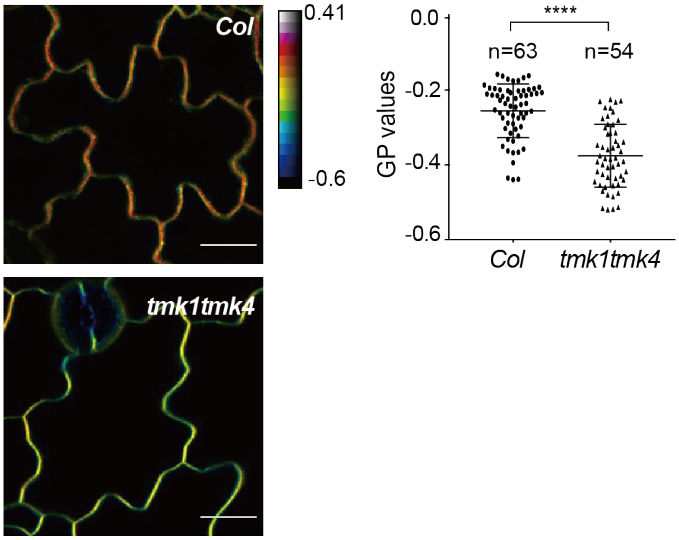

**fig S3. Compared to wild-type (*Col-0*), plasma membrane order is significantly reduced in the *tmk1tmk4* double mutant. (left panel)** Representative GP images of pavement cells in wild-type and *tmk1tmk4* double mutant obtained after di-4-ANEPPDHQ staining. **(right panel)** Quantitative analysis of mean GP value extracted from the plasma membrane of multiple pavement cells. Scale bars = 15 μm. *n* represents the number of independent cells. Data were presented as mean ± SD. See methods for detailed statistical analyses.

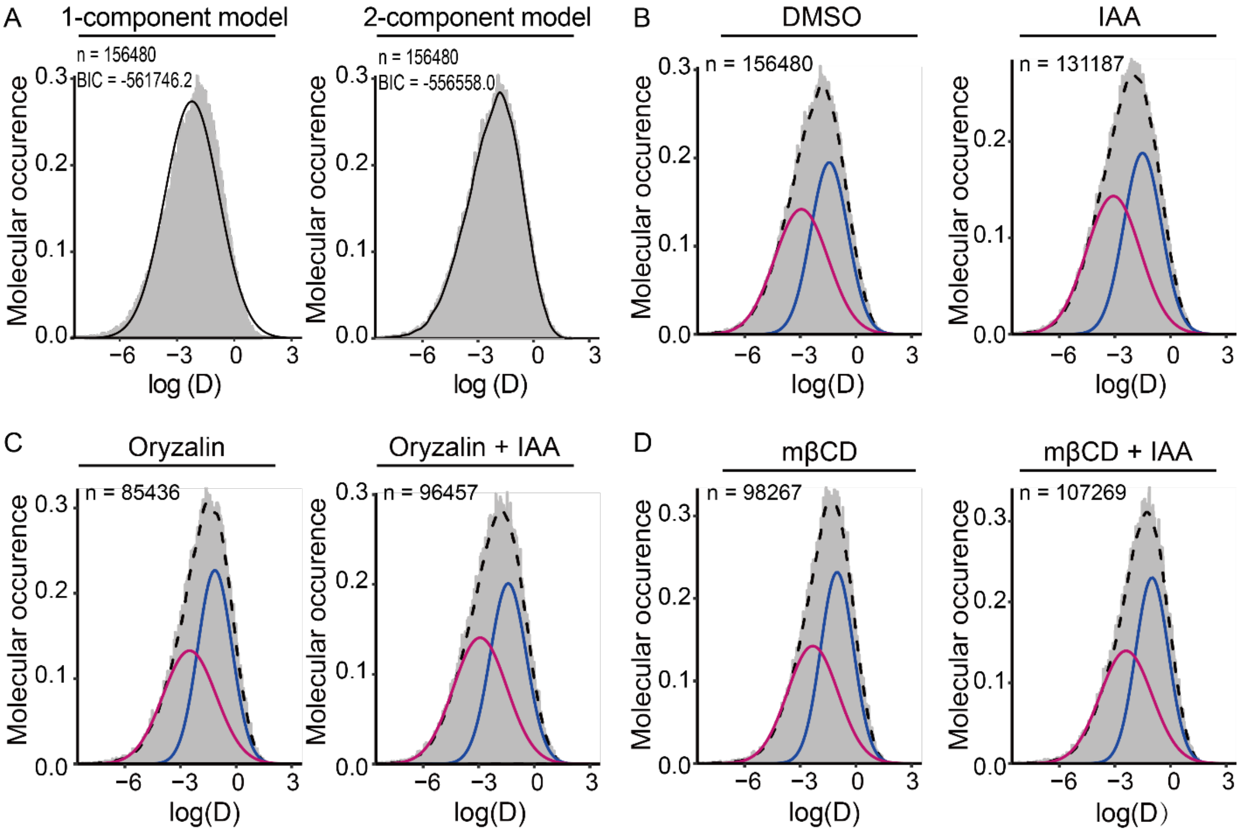

**fig S4. Raw data of diffusion coefficient of TMK1 particles obtained from single-particle tracking experiments. (A)** The density histogram of diffusion coefficients for TMK1 particles and the results of the fits to the one-component model (**left panel**) and two-component (**right panel**) model. The fitting of the density curve (solid line) to the density histogram (gray area) is significantly improved when the two-component mixture model is used. This is consistent with the higher Bayesian Information Criterion (BIC) values obtained from the two-component mixture model. **(B)** Raw data of diffusion coefficient of TMK1 particles in mock and IAA-treated conditions. **(C)** Raw data of diffusion coefficient of TMK1 particles in oryzalin- and oryzalin/IAA-treated conditions. **(D)** Raw data of diffusion coefficient of TMK1 particles in methyl-β- cyclodextrin (mβCD)- and mβCD /IAA- treated conditions. The gray area indicates the density histogram of diffusion coefficients obtained by a mean squared displacement analysis of single-particle trajectories. The pink and blue curves shown in the plot correspond to the individual Gaussian density components in the mixture distribute on, each scaled by the estimated probability of an observation being drawn from that component distribution. The heavy dashed line shows the nonparametric density estimate generated by the density function in the R mClust Package with the default settings. *n* represents the total number of particles.

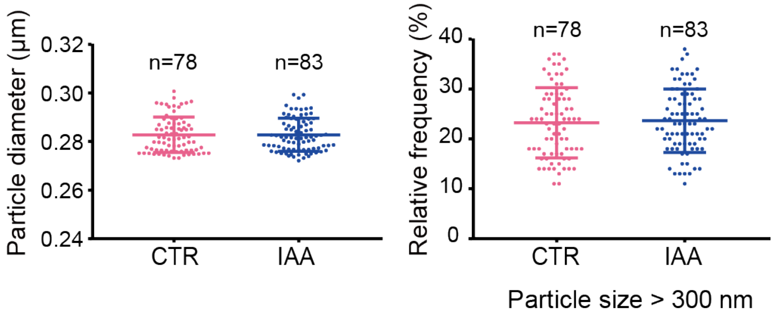

**fig S5. Auxin treatment (100nM, 10min) shows no significant effect on the particle size of the FERONIA (FER) receptor-like kinase at the plasma membrane of pavement cells.** Transgenic Arabidopsis expression a YFP-FER fusion protein driven by the native FER promoter in the *fer-4* mutant background was used in this experiment. **Left panel:** Quantitative comparison of FER-GFP particle size between control and IAA treatment. No statistical differences were observed between the control and treatment group. **Right panel:** Relative frequency of FER-GFP particles with size larger than 300nm between control and IAA treatment. No statistical differences were observed between the control and treatment group. *n* represents the number of independent cells.

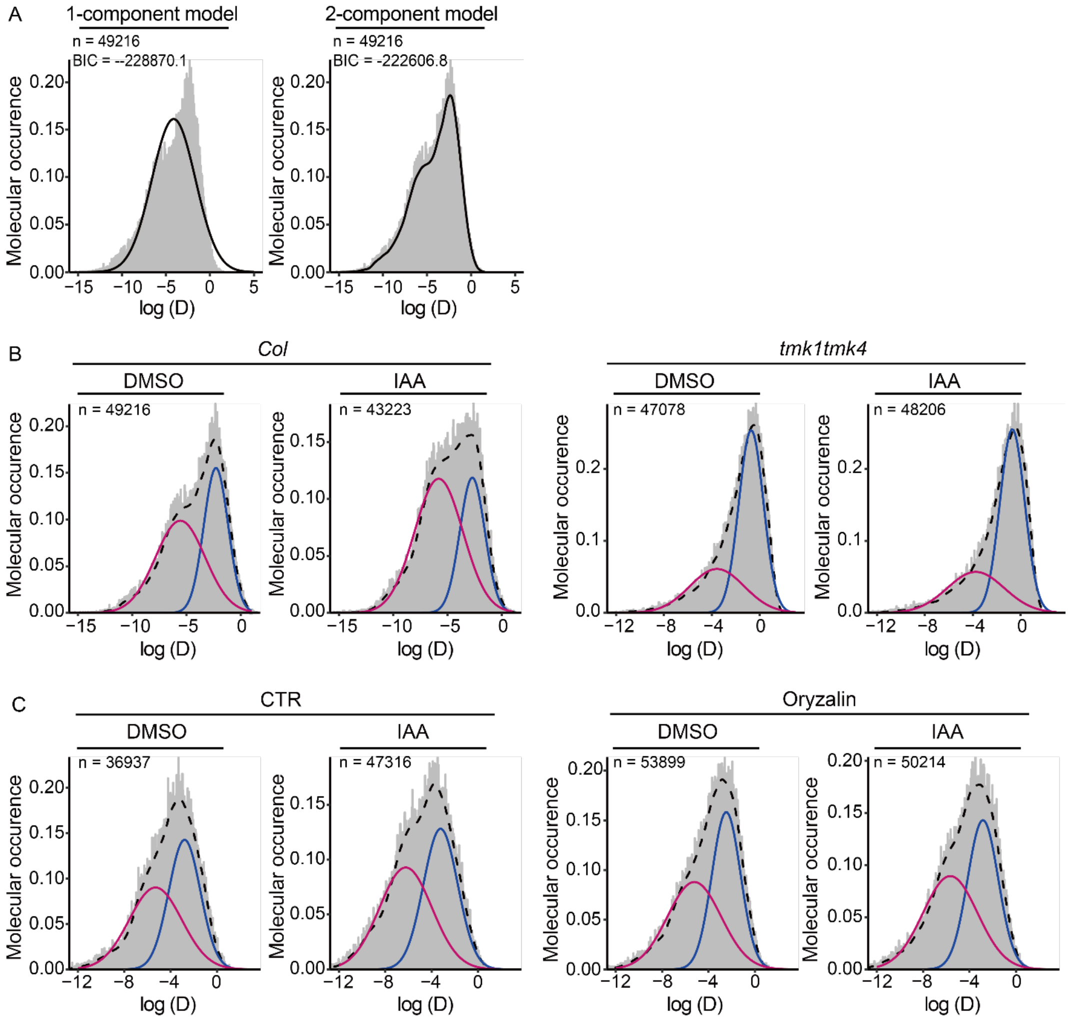

**fig S6. Raw data of diffusion coefficient of flotillin1 particles obtained from single-particle tracking experiments. (A)** The density histogram of diffusion coefficients for flotillin1 particles and the results of the fits to the one-component model (**left panel**) and two-component (**right panel**) model. The fitting of the density curve (solid line) to the density histogram (gray area) is significantly improved when the two-component mixture model is used. This is consistent with the higher Bayesian Information Criterion (BIC) values obtained from the two-component mixture model. **(B)** Raw data of diffusion coefficient of flotillin1 particles obtained from single-particle tracking experiments in mock and auxin-treated conditions in wild-type (*Col-0*) and *tmk1tmk4* double mutant background. **(C)** Raw data of diffusion coefficient of flotillin1 particles obtained from single-particle tracking experiments in mock, auxin-treated, and/or oryzalin-treated conditions. The gray area indicates the density histogram of diffusion coefficients obtained by a mean squared displacement analysis of single-particle trajectories. The pink and blue curves shown in the plot correspond to the individual Gaussian density components in the mixture distribution, each scaled by the estimated probability of an observation being drawn from that component distribution. The heavy dashed line shows the nonparametric density estimate generated by the density function in the R mClust Package with the default settings. *n* represents the total number of particles.

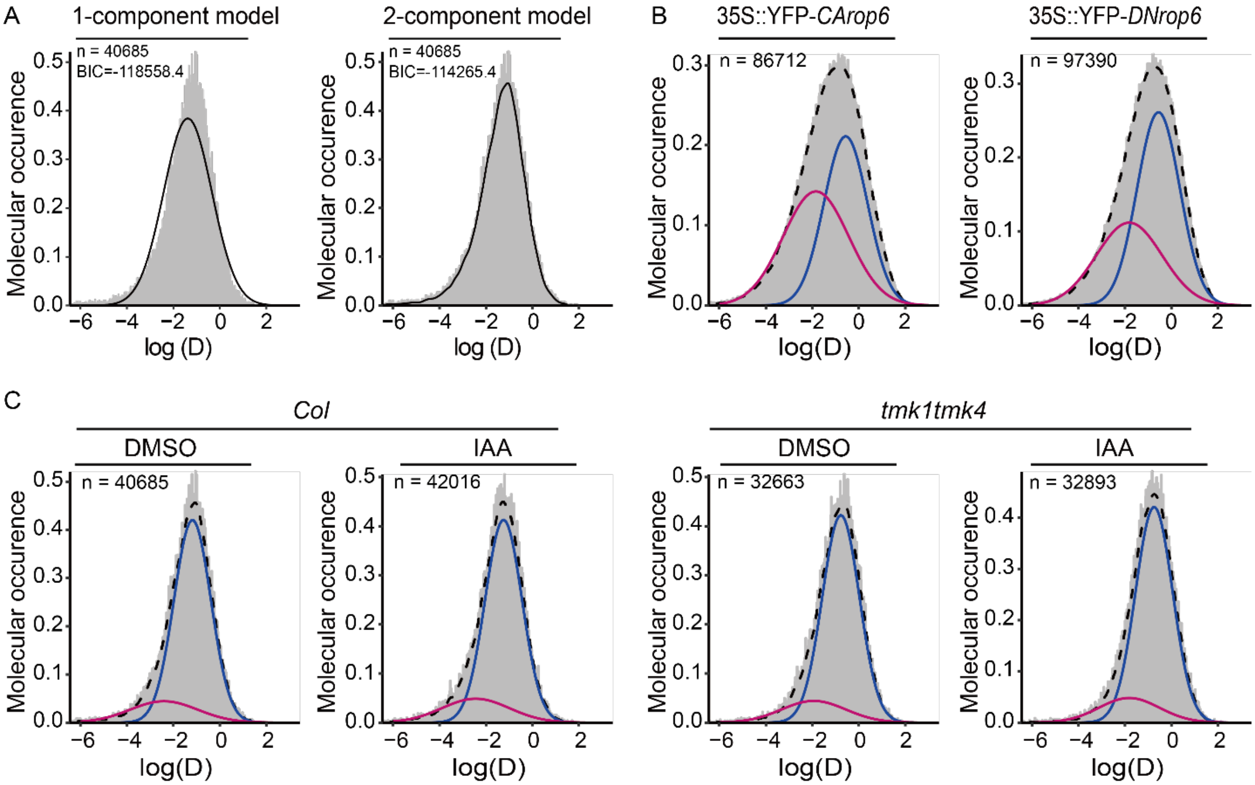

**fig S7. Raw data of diffusion coefficient of ROP6 particles obtained from single-particle tracking experiments. (A)** The density histogram of diffusion coefficients for ROP6 particles and the results of the fits to the one-component model (left panel) and two-component (right panel) model. The fitting of the density curve (solid line) to the density histogram (gray area) is significantly improved when the two-component mixture model is used. This is consistent with the higher Bayesian Information Criterion (BIC) values obtained from the two-component mixture model. **(B)** Raw data of diffusion coefficient of GFP-*CArop6* and GFP-*DNrop6* particles. **(C)** Raw data of diffusion coefficient of ROP6 particles with or without IAA treatment in wild-type (*Col-0*) or *tmk1tmk4* double mutant background. The gray area indicates the density histogram of diffusion coefficients obtained by a mean squared displacement analysis of single-particle trajectories. The pink and blue curves shown in the plot correspond to the individual Gaussian density components in the mixture distribution, each scaled by the estimated probability of an observation being drawn from that component distribution. The heavy dashed line shows the nonparametric density estimate generated by the density function in the R mClust Package with the default settings. *n* represents the total number of particles.

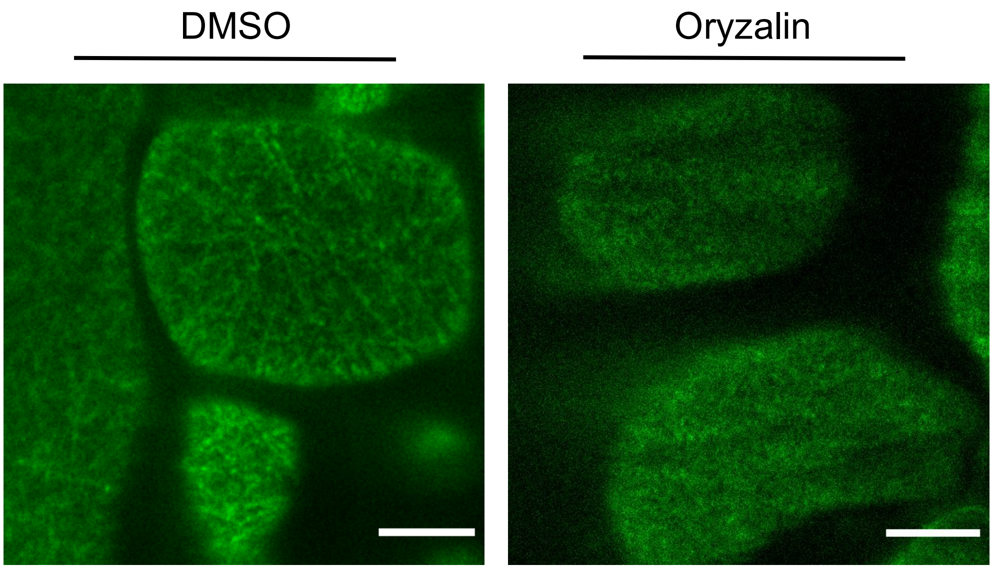

**fig S8. Depolymerization of microtubules by oryzalin treatment abolishes the accumulation of *CArop6* in filamentous structures.** Representative TIRF images of pavement cells of 2-3 days-old cotyledons expressing *35S:GFP-CArop6* without (**left panel**) or with oryzalin treatment (5µM, 2hrs) (**right panel**). Scale bars = 10 µm.

**A**

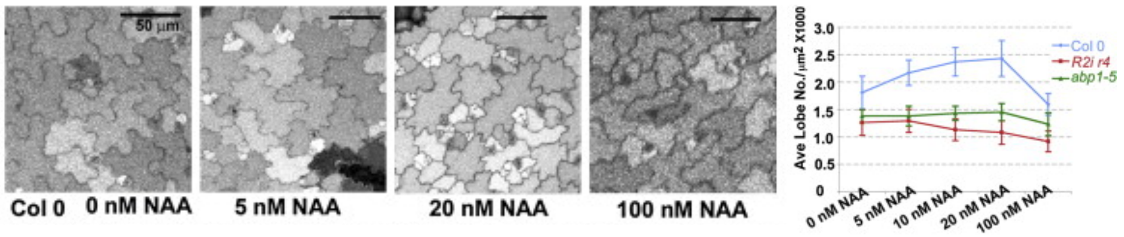

Xu, Tongda, et al. "Cell surface-and rho GTPase-based auxin signaling controls cellular interdigitation in Arabidopsis." Cell 143.1 (2010): 99-110.

**B**

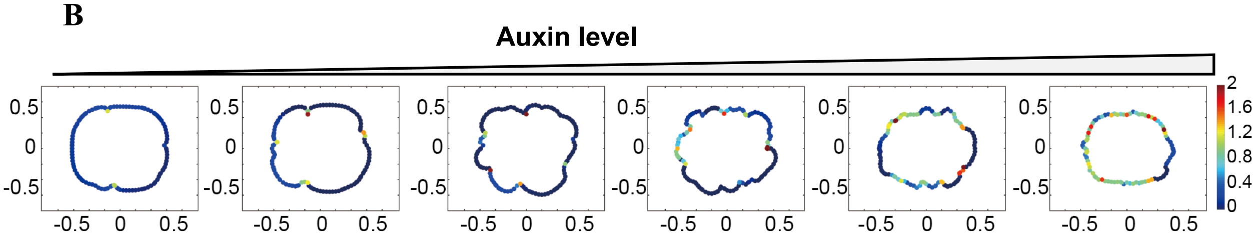

**fig S9. Simulation reproduced the dose-dependent effect of auxin on pavement cell (PC) interdigitation. (A)** Auxin (NAA) dosage responses in wild-type PCs. The data was previously published in (*3*). NAA promoted PC interdigitation upto 20nM and the promotion effect decreased with 100nM NAA. **(B)** Simulated PCs with altered auxin levels.

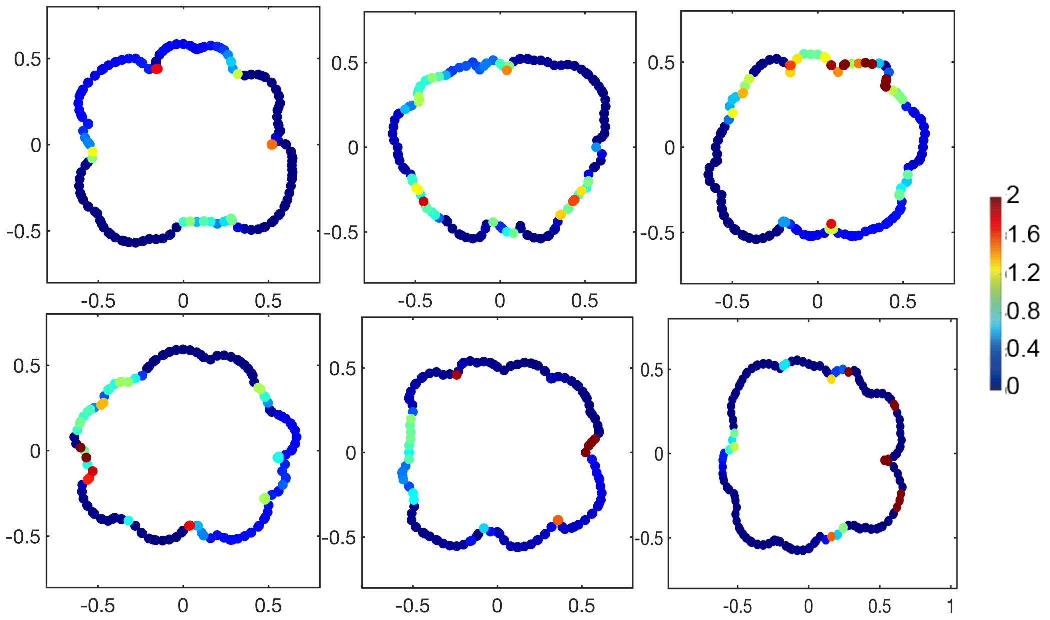

**fig S10. Simulated images of pavement cells generated using the settings for the mock treatment shown in Figure 6E**. The color scale indicates the level of active ROP6.

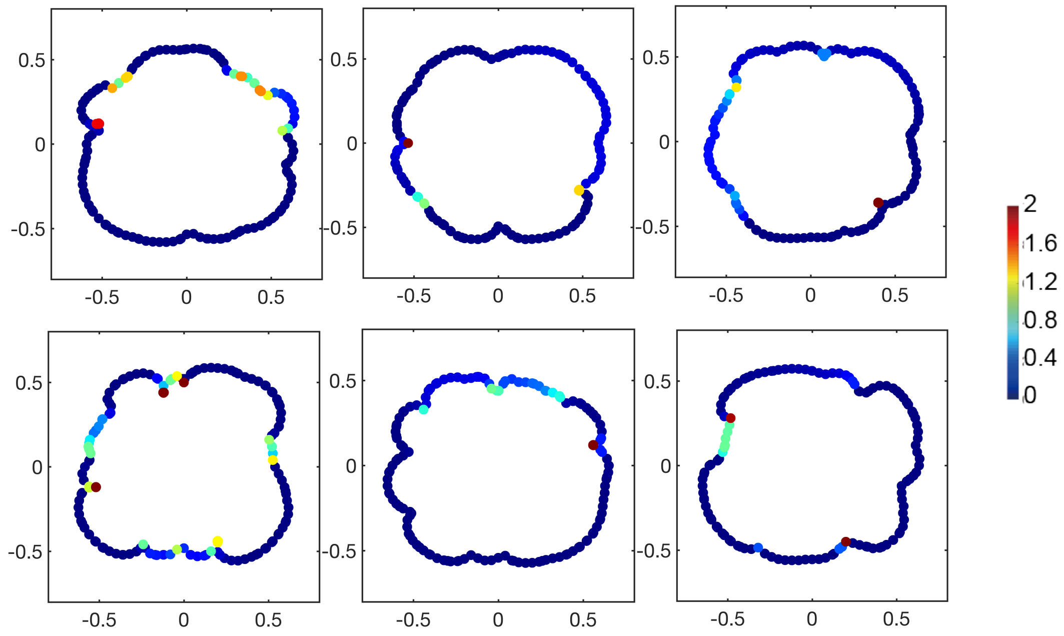

**fig S11. Simulated images of pavement cells generated using the settings for the mβCD treatment shown in Figure 6E**. The color scale indicates the level of active ROP6.

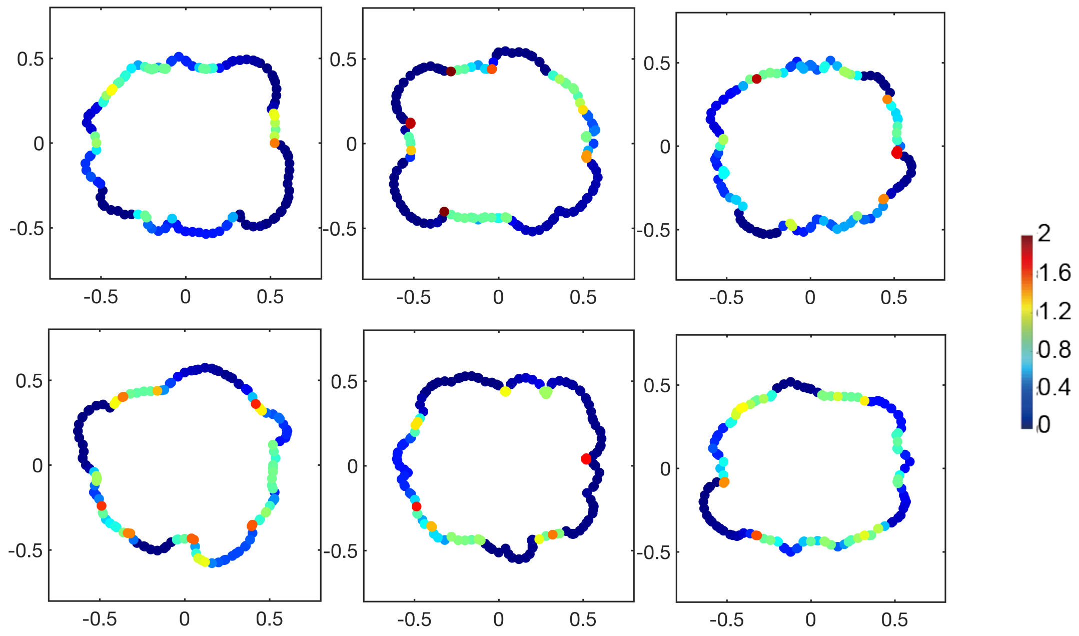

**fig S12. Simulated images of pavement cells generated using the settings for the auxin treatment shown in Figure 6E**. The color scale indicates the level of active ROP6.

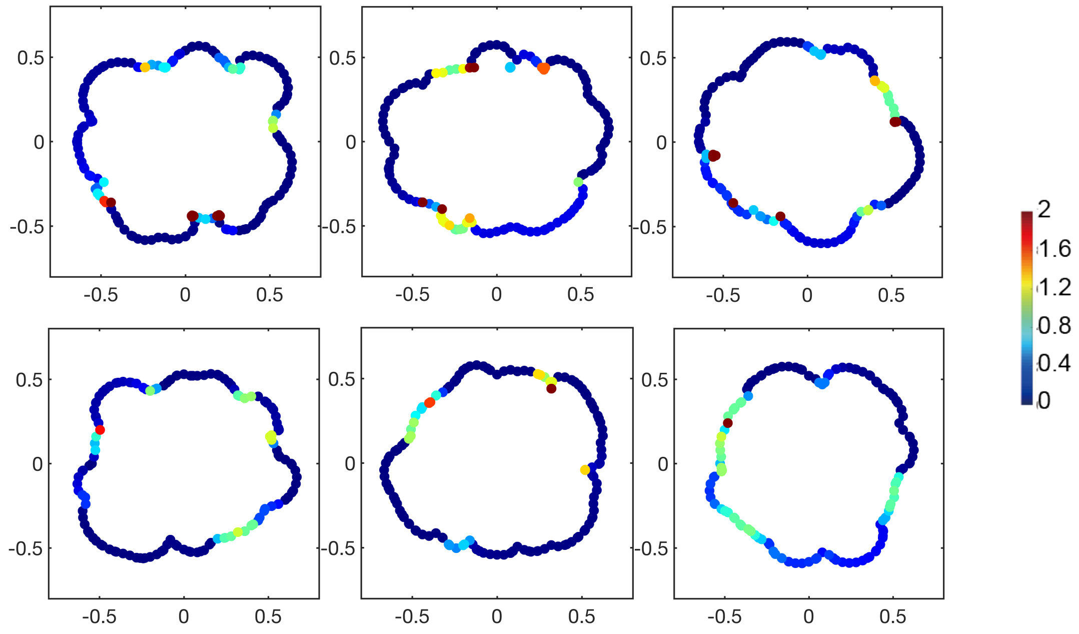

**fig S13. Simulated images of pavement cells generated using the settings for the combined auxin and mβCD treatment shown in Figure 6E**. The color scale indicates the level of active ROP6.

### References

1. Y. Fu, T. Xu, L. Zhu, M. Wen, Z. Yang, A ROP GTPase signaling pathway controls cortical microtubule ordering and cell expansion in Arabidopsis. *Current biology : CB* **19**, 1827-1832 (2009); published online EpubNov 17 (10.1016/j.cub.2009.08.052).
2. S. J. Clough, A. F. Bent, Floral dip: a simplified method for Agrobacterium-mediated transformation of Arabidopsis thaliana. *The Plant journal : for cell and molecular biology* **16**, 735-743 (1998); published online EpubDec (
3. T. Xu, M. Wen, S. Nagawa, Y. Fu, J. G. Chen, M. J. Wu, C. Perrot-Rechenmann, J. Friml, A. M. Jones, Z. Yang, Cell surface- and rho GTPase-based auxin signaling controls cellular interdigitation in Arabidopsis. *Cell* **143**, 99-110 (2010); published online EpubOct 1 (10.1016/j.cell.2010.09.003).
4. Y. Fu, Y. Gu, Z. Zheng, G. Wasteneys, Z. Yang, Arabidopsis interdigitating cell growth requires two antagonistic pathways with opposing action on cell morphogenesis. *Cell* **120**, 687-700 (2005); published online EpubMar 11 (10.1016/j.cell.2004.12.026).
5. D. Lin, L. Cao, Z. Zhou, L. Zhu, D. Ehrhardt, Z. Yang, Y. Fu, Rho GTPase signaling activates microtubule severing to promote microtubule ordering in Arabidopsis. *Current biology : CB* **23**, 290-297 (2013); published online EpubFeb 18 (10.1016/j.cub.2013.01.022).
6. T. Xu, N. Dai, J. Chen, S. Nagawa, M. Cao, H. Li, Z. Zhou, X. Chen, R. De Rycke, H. Rakusova, W. Wang, A. M. Jones, J. Friml, S. E. Patterson, A. B. Bleecker, Z. Yang, Cell surface ABP1-TMK auxin-sensing complex activates ROP GTPase signaling. *Science* **343**, 1025-1028 (2014); published online EpubFeb 28 (10.1126/science.1245125).
7. P. A. Hemsley, L. Taylor, C. S. Grierson, Assaying protein palmitoylation in plants. *Plant methods* **4**, 2 (2008); published online EpubJan 11 (10.1186/1746-4811-4-2).
8. D. Wessel, U. I. Flugge, A method for the quantitative recovery of protein in dilute solution in the presence of detergents and lipids. *Analytical biochemistry* **138**, 141-143 (1984); published online EpubApr (
9. P. Gerbeau-Pissot, C. Der, M. Grebe, T. Stanislas, Ratiometric Fluorescence Live Imaging Analysis of Membrane Lipid Order in Arabidopsis Mitotic Cells Using a Lipid Order-Sensitive Probe. *Methods in molecular biology* **1370**, 227-239 (2016)10.1007/978-1-4939-3142-2_17).
10. X. Zhao, R. Li, C. Lu, F. Baluska, Y. Wan, Di-4-ANEPPDHQ, a fluorescent probe for the visualisation of membrane microdomains in living Arabidopsis thaliana cells. *Plant physiology and biochemistry : PPB / Societe francaise de physiologie vegetale* **87**, 53-60 (2015); published online EpubFeb (10.1016/j.plaphy.2014.12.015).
11. D. M. Owen, C. Rentero, A. Magenau, A. Abu-Siniyeh, K. Gaus, Quantitative imaging of membrane lipid order in cells and organisms. *Nature protocols* **7**, 24-35 (2012); published online EpubJan (10.1038/nprot.2011.419).
12. Y. Fu, H. Li, Z. Yang, The ROP2 GTPase controls the formation of cortical fine F-actin and the early phase of directional cell expansion during Arabidopsis organogenesis. *The Plant cell* **14**, 777-794 (2002); published online EpubApr (
13. M. Goulian, S. M. Simon, Tracking single proteins within cells. *Biophysical journal* **79**, 2188-2198 (2000); published online EpubOct (10.1016/S0006-3495(00)76467-8).
14. S. Osher, Sethian, J.A., Fronts propagating with curvature-dependent speed: algorithms based on Hamilton-Jacobi formulations. *Journal of Computational Physics* **79**, 12-49 (1988).
15. J. J. Xu, H. K. Zhao, An Eulerian formulation for solving partial differential equations along a moving interface. *J. Sci. Comput.* **19**, 573 (2003).

16. M. Souter, J. Topping, M. Pullen, J. Friml, K. Palme, R. Hackett, D. Grierson, K. Lindsey, hydra Mutants of Arabidopsis are defective in sterol profiles and auxin and ethylene signaling. *The Plant cell* **14**, 1017-1031 (2002); published online EpubMay (
17. S. D. Gilk, D. C. Cockrell, C. Luterbach, B. Hansen, L. A. Knodler, J. A. Ibarra, O. Steele-Mortimer, R. A. Heinzen, Bacterial colonization of host cells in the absence of cholesterol. *PLoS pathogens* **9**, e1003107 (2013); published online EpubJan (10.1371/journal.ppat.1003107).
18. J. C. Jang, S. Fujioka, M. Tasaka, H. Seto, S. Takatsuto, A. Ishii, M. Aida, S. Yoshida, J. Sheen, A critical role of sterols in embryonic patterning and meristem programming revealed by the fackel mutants of Arabidopsis thaliana. *Genes & development* **14**, 1485-1497 (2000); published online EpubJun 15 (
19. S. Men, Y. Boutte, Y. Ikeda, X. Li, K. Palme, Y. D. Stierhof, M. A. Hartmann, T. Moritz, M. Grebe, Sterol-dependent endocytosis mediates post-cytokinetic acquisition of PIN2 auxin efflux carrier polarity. *Nature cell biology* **10**, 237-244 (2008); published online EpubFeb (10.1038/ncb1686).
